# Predatory orienting is jointly controlled by optic tectum and pretectum

**DOI:** 10.1101/2025.11.10.687578

**Authors:** Paride Antinucci, Susana Colinas-Fischer, Isaac H. Bianco

## Abstract

The optic tectum and pretectum have been implicated in hunting behaviour in several species but how activity in these regions is coordinated to control target-directed orienting manoeuvres is not understood. We performed two-photon calcium imaging across these brain regions in larval zebrafish, which revealed space and rate-coded premotor activity associated with graded, prey-directed eye and tail movements. Consistent with imaging data, intensity-modulated optogenetic stimulation of pretectum induced progressively more lateralised contraversive orienting behaviour and spatially patterned stimulation of optic tectum revealed a motor map along the anterior-posterior axis that primarily generated ipsiversive responses. Anatomical tracing and laser axotomies indicated that pretectal neurons control contraversive turns via a crossed pretectobulbar pathway whereas optic tectum shows topographically patterned output with a major uncrossed projection to hindbrain. Our data support a model in which premotor activity across pretectum and optic tectum collectively controls predatory orienting via both crossed and uncrossed (pre)-tectobulbar pathways.

## Introduction

Hunting is a conserved behaviour, which often requires predators to generate a succession of precise, target-directed orienting movements to pursue and capture moving prey. How is predatory orienting controlled? The midbrain superior colliculus (SC; or optic tectum (OT) in non-mammalian vertebrates) has a well-established role in controlling orienting behaviours, including in the context of hunting^1^. Canonical models of SC/OT function emphasise a ‘space code’, wherein topographically organised sensory input is transformed to spatially localised premotor activity, which specifies the amplitude (or end-point) of the orienting movement^2^. However, although this model has substantial support, there are a number of open questions. First, many studies of SC/OT have been in the context of localisation responses to transient visual cues, but this behaviour might have distinct neural control as compared to hunting-related pursuit^3,4^ (and see Discussion). Second, much of our understanding derives from single-unit recordings and it is unclear how orienting behaviours are encoded by population activity across SC/OT, including across the two brain hemispheres. Third, midbrain commands are conveyed to tegmental motor generators responsible for the eye, head, and body movements that comprise the orienting response and while much evidence suggests the key efferent pathway is the crossed tectobulbar tract^1^, some studies support a role for the uncrossed descending pathway^5,6^. Fourth, some aspects of orienting responses are preserved after tectal inactivation and it is not clear how SC/OT functions in parallel with other brain regions to generate target-directed behaviour^2^.

The larval zebrafish is a powerful model to address such questions. Its hunting behaviour has been extensively characterised^7–13^ and can be evoked in tethered animals amenable to cellular-resolution functional imaging and other circuit interrogation methods^9,14^. Hunting is primarily visually guided, depending on retinal input to OT^15^, which contains several classes of neuron that encode prey-like visual features^6,14,16–19^. In line with the conserved role of SC/OT in predatory orienting, spatially localised bursts of tectal activity predict the occurrence and direction of hunting responses ^14,20^ and hunting outputs can be evoked by stimulation of anterior-ventral OT^21^. However, in addition to OT, recent studies have also cemented a role for the pretectum in hunting control, specifically pretectal nuclei adjacent to retinal arborisation field 7 (hereafter Pt7). Arborisation field 7 is innervated by retinal ganglion cells that are tuned to prey-like visual features^22^ and a subset of Pt7 cells, labelled by the *u508* transgene, function as a command system to evoke hunting behaviour^18^. Moreover, the observation that optogenetic stimulation of *u508* neurons evokes hunting responses with consistent laterality suggests that Pt7 might also influence the directionality of hunting behaviour, perhaps operating alongside OT to steer predatory actions towards prey targets.

Here, we set out to dissect the roles of Pt7 and OT in predatory orienting. Using cellular-resolution calcium imaging during naturalistic hunting, we observed that target-directed responses were associated with distributed premotor activity across Pt7 and OT. This included activity in anterior regions contralateral to the direction of orienting movements, as well as an unexpected locus of activity in posterior OT, ipsilateral to turn direction. The amplitude of eye and body rotations covaried with the spatial distribution of premotor activity as well as the level of activity of individual cells, pointing to a role for both space and rate-coding. Optogenetic stimulation supported a causal role for these components of Pt7-OT activity and circuit tracing and laser axotomies revealed a correspondence between functional data and the topography of descending projections to ipsilateral and contralateral hindbrain. In sum, our data are consistent with a joint role for Pt7 and OT, and both crossed and uncrossed (pre)-tectobulbar pathways, in controlling the precise, target-directed orienting responses that are essential for hunting.

## Results

### Topography of hunting-specific sensory and premotor activity across Pt7-OT

Here, we set out to understand the neural basis of target-directed eye and body movements during hunting. First, we examined how visual sensory representations of prey relate to premotor activity that directs hunting actions by combining a virtual hunting assay with 2-photon calcium imaging in tethered larval zebrafish^9,14^.

Zebrafish generated predatory orienting movements, the magnitude and direction of which scaled with prey location. We presented larvae (6–7 days post-fertilisation (dpf)) with small, prey-like moving spots, which swept horizontally across the frontal visual field [Figure 1A], and specifically identified hunting responses by detecting convergent saccades, which are used by larval zebrafish to initiate hunting routines in coordination with orienting swims ^9,23^. Larvae responded to visual targets within a ∼ 120*^◦^* reactive perceptive field centred on their extended midsagittal plane (midline) [Figure 1B], similar to freely swimming animals hunting live prey^9,11,13,24,25^. Within this region, eye and tail movements varied in direct proportion to prey azimuth. Specifically, tail bend angle [Figure 1C], as well as the conjugate component of post-saccadic gaze (hereafter ‘gaze angle’) [Figure 1D], scaled linearly with target eccentricity. Thus, in accordance with natural hunting^10,11,23^, tethered larvae generate smooth, continuous adjustments in their eye and tail movements to orient towards prey targets.

**Figure 1:**
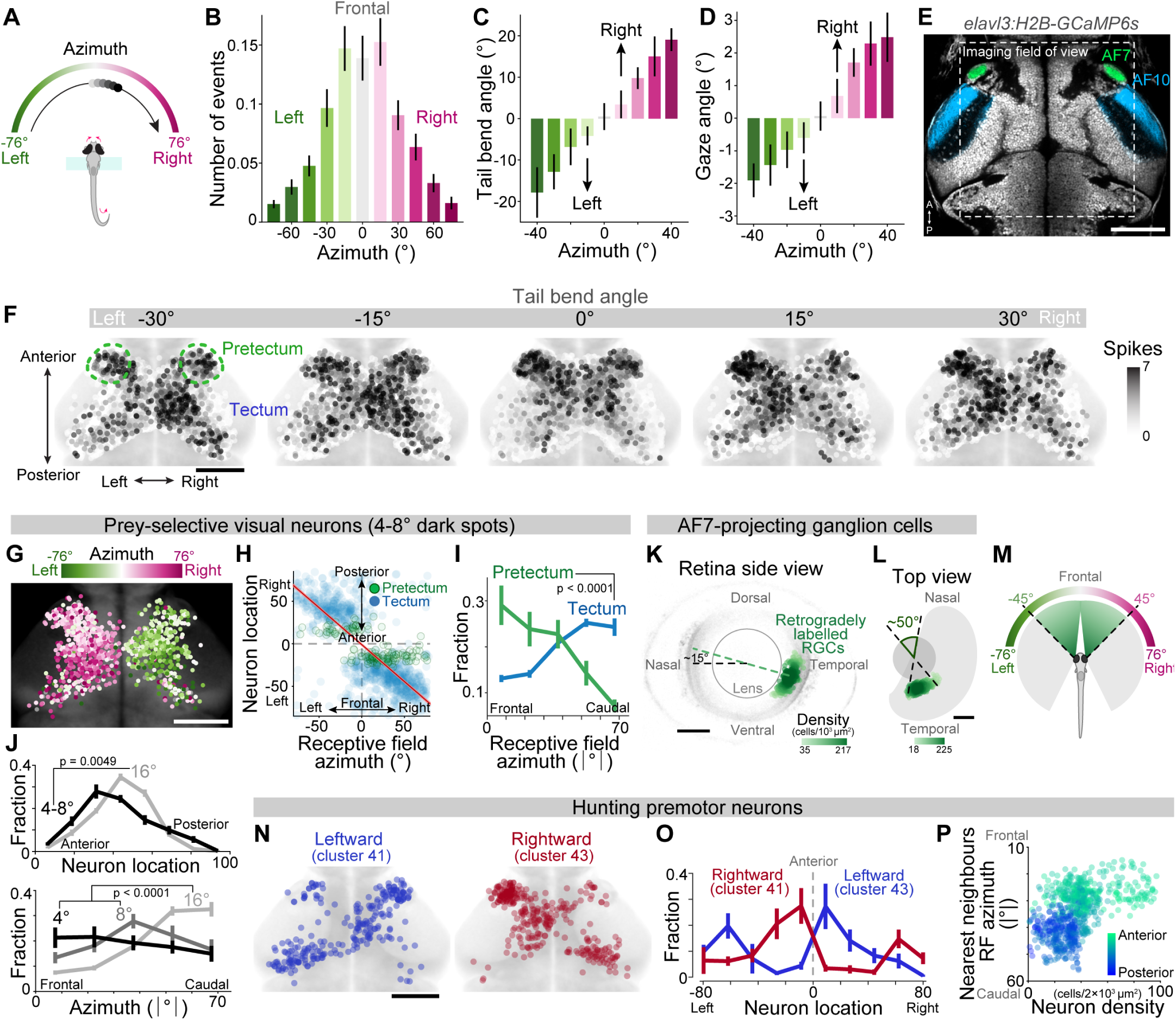
Predatory orienting and associated sensorimotor activity in Pt7-OT. (A) Schematic of virtual hunting assay. (B) Occurrence of hunting responses (convergent saccades) as a function of target azimuth. Mean ± SEM were computed across animals, as for all other plots (*N* = 96 larvae producing 1,424 responses). (C) Tail bend angle versus target azimuth. (D) Gaze angle (mean of left and right post-saccadic eye position) versus target azimuth. (E) Imaging field of view (310 × 310 *µ*m), superimposed on Tg(*elavl3*:H2B-GCaMP6s) expression (grey). Retinal innervation of pretectal AF7 (green) and tectal AF10 (blue) is visualised using the Tg(*atoh7*:gapRFP) transgene. Scale bar: 100*µ*m. (F) Activity maps for hunting events binned by tail bend angle. Each neuron is colour-coded according to the mean number of OASIS-inferred spikes across events in the corresponding bin (25,802 neurons from 28 larvae). Scale bar: 100 *µ*m. (G) Retinotopic mapping of visual azimuth. Prey-selective visual neurons (clusters 1–7, 1,106 cells) colour-coded by location of RF centre. (H) Soma location (normalised position along anterior-posterior axis) versus RF location for prey-selective visual neurons (clusters 1–7). Global retinotopy indicated by linear regression fit (red lines with 95% CI shading) for all spot-selective cells (clusters 1–31). (I) Pt7 neurons over-represent frontal visual space. Distribution of absolute RF location for spot-selective cells in Pt7 and OT. *p*-value from two-sample Kolmogorov-Smirnov test. (J) Soma locations (top) and RF location (bottom) for neurons tuned to prey-like moving spots of different sizes. *p*-values from two-sample Kolmogorov-Smirnov test. (K,L) Retrograde labelling of RGCs innervating AF7. Somata shown in saggital (K) and dorsal (L) views of the retina, coded by local 2D cell density. Green dashed line in (K) passes through centre of mass of AF7-projecting RGCs and centre of lens, black dashed line indicates horizontal retinal axis. Dashed lines in (L) indicate angle subtended in the horizontal plane that is viewed by these RGCs. Scale bar: 50 *µ*m. (M) Estimated visual field viewed by AF7-projecting RGCs. Resting eye position was taken as 14.2 ± 3.8^◦^ (*N* = 192 eyes) with each eye assumed to view 163*^◦^*in the horizontal plane ^76^. (N) Maps of direction-selective hunting premotor neurons (cluster 41, blue, 273 cells; cluster 43, red, 357 cells). Scale bar: 100 *µ*m. (O) Distribution of hunting premotor neurons along anterior-posterior axis. Mean ± SEM across *N* = 22 animals. (P) Local 2D density of premotor neurons versus mean absolute RF centre of 20-nearest neighbour prey-selective neurons. Each point represents a premotor neuron colour-coded according to its anterior-posterior location.

To understand how these target-directed actions might be controlled by tectal and pretectal populations, we performed calcium imaging of Tg(*elavl3*:H2B-GCaMP6s) larvae. Our imaging volume encompassed the region of pretectum surrounding arborisation field 7 (i.e. Pt7) as well as cells in the periventricular layer of OT, and spanned both brain hemispheres [Figure 1E]. Individual neurons were automatically segmented and brain registration was used to map their locations to a standard coordinate space (ZBB brain atlas^26,27^). To account for indicator dynamics, we used the OASIS^28^ deconvolution algorithm to estimate a ‘spiking’ process for each cell. Hereafter, we refer to this representation of activity as inferred spikes (spk), while recognising that it does not equate to action potential counts or firing rates. Using our full set of imaged neurons, we generated activity maps spanning Pt7-OT, binned according to the amplitude of predatory orienting movements [Figure 1F]. This revealed that forward-directed hunting actions (i.e. 0*^◦^* bin) were associated with bilaterally symmetric activity in anterior OT and Pt7, whereas lateralised actions were associated with elevated activity in the brain hemisphere contralateral to the direction of tail and gaze displacement (i.e. left Pt7-OT activity associated with rightwards hunting responses).

Next, we sought to decompose these activity maps to distinguish sensory representations of prey from premotor activity that might be responsible for directing hunting actions. To do this, we built linear encoding models for every neuron and clustered the resulting visuomotor tuning vectors (VMVs) to identify groups of cells with distinct functional properties (Methods and^18^): Several clusters encoded specific visual cues (e.g. moving spots, looming stimuli, luminance changes) while the activity of other neurons was explained by behavioural variables (e.g. eye and tail kinematics) [Figure S1]. For spot-selective neurons (clusters 1–31), we estimated each cells’ spatial receptive field (RF) by relating the time-course of inferred spikes to the trajectory of the visual target. As expected, this revealed a retinotopic representation of visual azimuth along the anterior-posterior axis of Pt7-OT such that frontal space was represented in Pt7 and anterior OT (aOT), whilst more eccentric locations mapped to medial (mOT), and posterior tectum (pOT) [Figure 1G-I]. Neurons that were most selective for prey-like visual features (tuned to small moving spots, 4–8*^◦^*) were particularly enriched in anterior locations with RFs viewing frontal visual space [Figure 1J]. The anterior RFs of Pt7 cells [Figure 1I] were explained by the retinal origin of ganglion cell inputs to this brain region. We showed this by photoactivating PA-GFP in retinorecipient AF7, which resulted in a high density of retrogradely labelled retinal ganglion cells in the ventral-temporal retina [Figure 1K,L], compatible with single-RGC morphology data^29^ and corresponding to the location of the retinal high acuity area (HAA, a.k.a ‘fovea’) ^30,31^. We estimated that at resting ocular vergence, this region views ∼ 90*^◦^* of frontal visual space [Figure 1M], which corresponds well to the reactive perceptive field of the animal [c.f. Figure 1B] as well as similar estimates based on photoreceptor density^31^.

Next, we examined neurons with premotor activity encoding the expression of hunting actions. In accordance with our previous study^18^, several clusters were specifically active when larvae initiated hunting with a convergent saccade paired with leftwards (cluster 41), rightwards (cluster 43) or either direction (cluster 42) of tail movement [Figure S1]. These neurons exclusively encoded hunting-specific orienting behaviour and were otherwise unresponsive to tail movements or prey-like visual cues (estimated from activity during non-response trials) [Figure S2]. We therefore refer to these cells as hunting premotor neurons. Directionally tuned premotor clusters had lateralised anatomical distributions [Figure 1N,O], which included a high density of cells in Pt7 and aOT contralateral to their preferred direction of movement. As such, in anterior locations, premotor encoding aligned with sensory encoding of prey-like stimuli in the contralateral, frontal visual field [Figure 1P]. Thus, Pt7-aOT in the right hemisphere, represented prey-like stimuli in the left frontal visual field and was enriched with premotor neurons tuned to leftwards (i.e. contraversive) hunting actions, and *vice versa* for the left hemisphere. However, in surprising contrast to this alignment of visual and motor maps in anterior regions, pOT was enriched with premotor cells tuned to *ipsiversive* orienting movements [Figure 1N].

In summary, neurons tuned to prey-like visual features are enriched in anterior Pt7-OT, in spatial register with premotor neurons encoding contraversive hunting movements. Yet, posterior tectum contains an additional cluster of premotor cells with ipsiversive tuning. Next, we aimed to understand how continuous variability in eye and tail kinematics relates to premotor activity.

### Predatory orienting is encoded by rate and space-coded activity in Pt7-OT

To assess how Pt7-OT encodes predatory orienting, we related the magnitude and spatial organisation of hunting premotor activity to the kinematics of eye and tail movements.

The activity of direction-selective premotor neurons progressively increased as zebrafish generated larger amplitude eye and tail movements in the cells’ preferred direction. We showed this by generating activity maps for cells belonging to cluster 41 (left-tuned) and cluster 43 (right-tuned) as a function of tail and eye movements organised into different amplitude bins [Figure 2A; Figure S3A]. Activity of cluster 41 neurons progressively increased as larvae initiated hunting with more negative (leftwards) tail angles [Figure 2A] and leftward-directed convergent saccades [Figure S3A], and *vice versa* for cells belonging to right-tuned cluster 43. Moreover, these activity maps revealed pronounced lateralisation in the patterns of premotor activity. Specifically, stronger orienting behaviour was associated with elevated activity of Pt7 and aOT in the contralateral hemisphere (i.e. in right Pt7-aOT for cluster 41 neurons as larvae oriented more strongly to the left). By contrast, in pOT, activity was elevated ipsilateral to turn direction. When zebrafish initiated hunting with a forward-directed swim, we observed an intermediate level of mirror-symmetric activity in cluster 41 and 43 that collectively resembled the symmetric pattern of Pt7-OT activity triggered on forward-directed hunting events [compare 0*^◦^* bin in Figure 2A to Figure 1F]. Thus, symmetric premotor activity encodes forward-directed predatory swims whereas lateralised hunting responses are associated with lateralised activation of directionally tuned premotor neurons.

**Figure 2:**
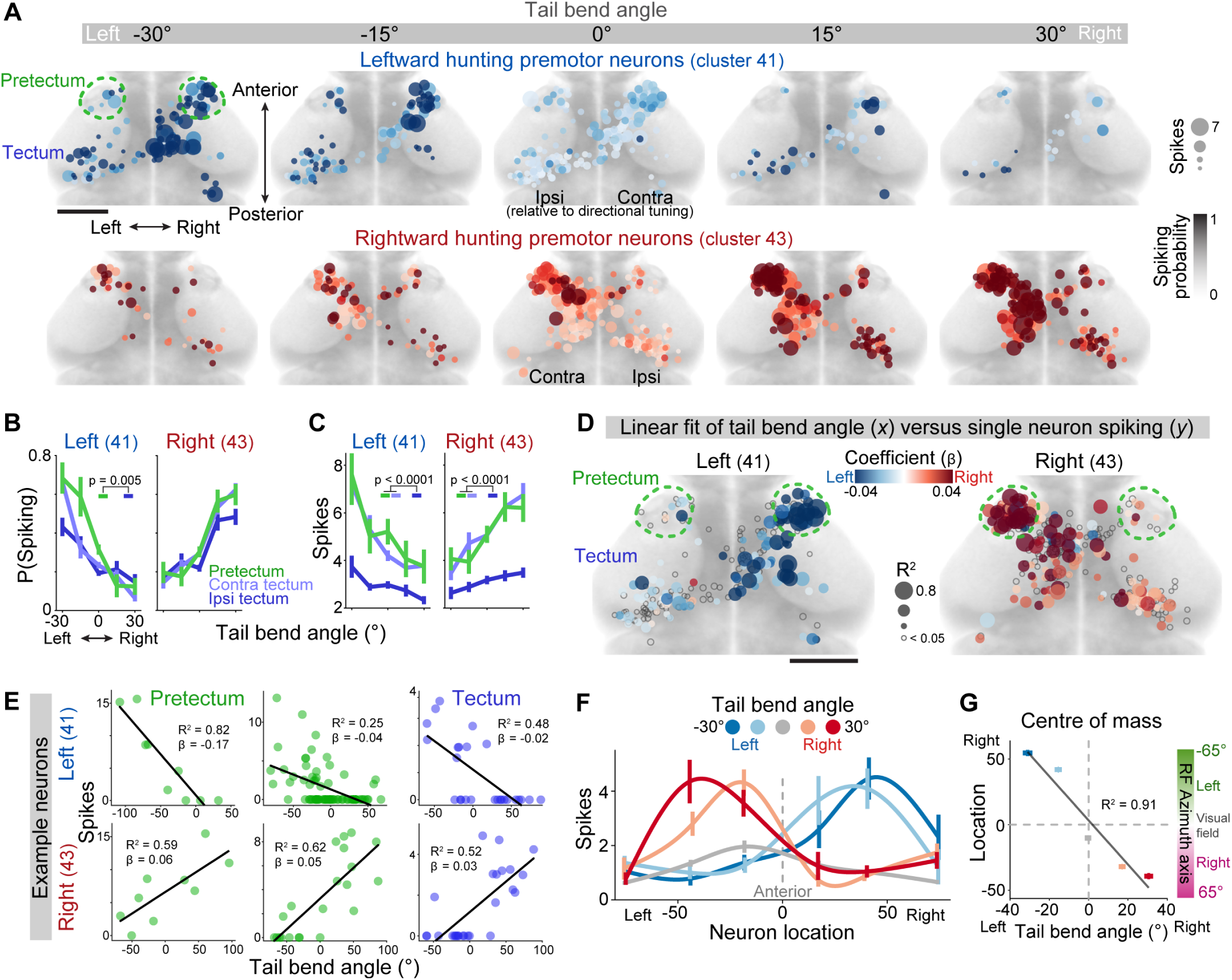
Rate- and space-coding of predatory orienting movements. (A) Activity maps for direction-selective premotor neurons (cluster 41, left-selective, *N* = 273 neurons; cluster 43, right-selective, *N* = 357 neurons from 22 larvae). OASIS-inferred spike probability (colour intensity) and spike counts (dot size) are shown for hunting manoeuvres binned by tail bend angle (bin centres indicated at top). (B,C) Spike probability (B) and spike counts (C) versus tail bend angle for premotor neurons in Pt7 (green), contralateral OT (light blue), and ipsilateral OT (dark blue), where ipsi/contralateral is with respect to the cells’ directional tuning. Mean ± SEM were computed across neurons, as for all the remaining plots (cluster 41: *N* = 39 pretectal neurons, *N* = 117 contra tectum, *N* = 117 ipsi tectum; cluster 43: *N* = 66 pretectal neurons, *N* = 183 contra tectum, *N* = 108 ipsi tectum). Tukey HSD-adjusted *p*-values for multiple comparison tests following two-way ANOVA. (D) Linear regression models relating single-cell spike counts (y) to tail bend angle (x). Fit coefficients (*β*, colour) and goodness-of-fit values (*R*^2^, dot size). Left angles are negative, so neurons encoding leftward movement have negative coefficients. (E) Examples of single neuron linear regression fits. (F) Spatial distribution of premotor activity along the anterior-posterior axis of Pt7-OT. Each trace represents a different bin of tail bend angle and shows spike counts across neurons (cluster 41 and 43 combined) binned by anatomical location. (G) Centre-of-mass of premotor activity for hunting manoeuvres binned by tail bend angle. The azimuth axis obtained from the linear regression fit in Figure 1H is also reported on the right.

Inspection of activity maps suggested predatory orienting was associated with both rate- and space-coded premotor activity. In support of rate-coding, we found that the mean number and probability of inferred spikes increased monotonically with the amplitude of tail and eye movements [Figure 2B,C, Figure S3B,C]. This modulation of population activity was evident in both Pt7 and OT. Furthermore, rate-coding was apparent at the level of individual neurons. We showed this by building single-cell linear regression models, which revealed that many neurons, especially in pretectum, positively scaled their activity as a function of predatory orienting, as indicated by their large slope coefficients and goodness-of-fit values [Figure 2D,E, Figure S3D,E].

Evidence from numerous species supports the existence of a motor map in SC/OT wherein the locus of tectal activity encodes the amplitude or end point of orienting movements^2^. Consistent with space coding, we found that the centre-of-mass of tectal population spiking shifted from anterior towards posterior OT with increasing (contralateral) steering angle [Figure 2F,G and Figure S3F,G].

Previously, we showed that Pt7 neurons with hunting premotor activity function as a command system to induce hunting^18^. Therefore, we examined the relationship between premotor activity and the probability of initiating hunting (as reported by saccadic eye convergence) and found that increasing activity of premotor clusters was associated with an increase in the probability of hunting [Figure S4]. This rate-code for hunting induction was also true at the single-neuron level [Figure S4C] and apparent for non-direction-selective premotor cells (cluster 42) [Figure S4D-F].

In sum, rate- and space-coded activity of hunting premotor neurons across Pt7-OT encodes the induction and steering of hunting behaviour. Whilst symmetric activity is associated with forward-directed hunting manoeuvres, imbalanced activity of directionally tuned populations encodes lateralised eye and tail movements. Premotor activity displayed pronounced topography wherein Pt7 and aOT were associated with contraversive predatory orienting whereas a locus of activity in pOT was recruited during ipsiversive orienting. To test the contribution of each component of these activity patterns to behaviour, we sought to use optogenetics to artificially impose activity on Pt7-OT circuits.

### The ***u523*** transgenic line preferentially labels Pt7-OT premotor neurons

To drive expression of optogenetic actuators, we screened for transgenic lines labelling cells in Pt7 and OT and identified two transgenes, *u523*:KalTA4 (hereafter *u523*) and *u524*:GAL4FF (hereafter *u524*) [Figure 3A]. By comparing the abundance of transgenically labelled neurons to a ‘pan-neuronal’ line (*elavl3*:H2B-GCaMP6s), we observed that *u523* preferentially labelled cells in anterior regions, Pt7 and aOT, while *u524* tended to label cells in more posterior tectum [Figure 3B,C; *N* = 11 *u523* larvae; *N* = 5 *u524*; 6–7 dpf].

**Figure 3:**
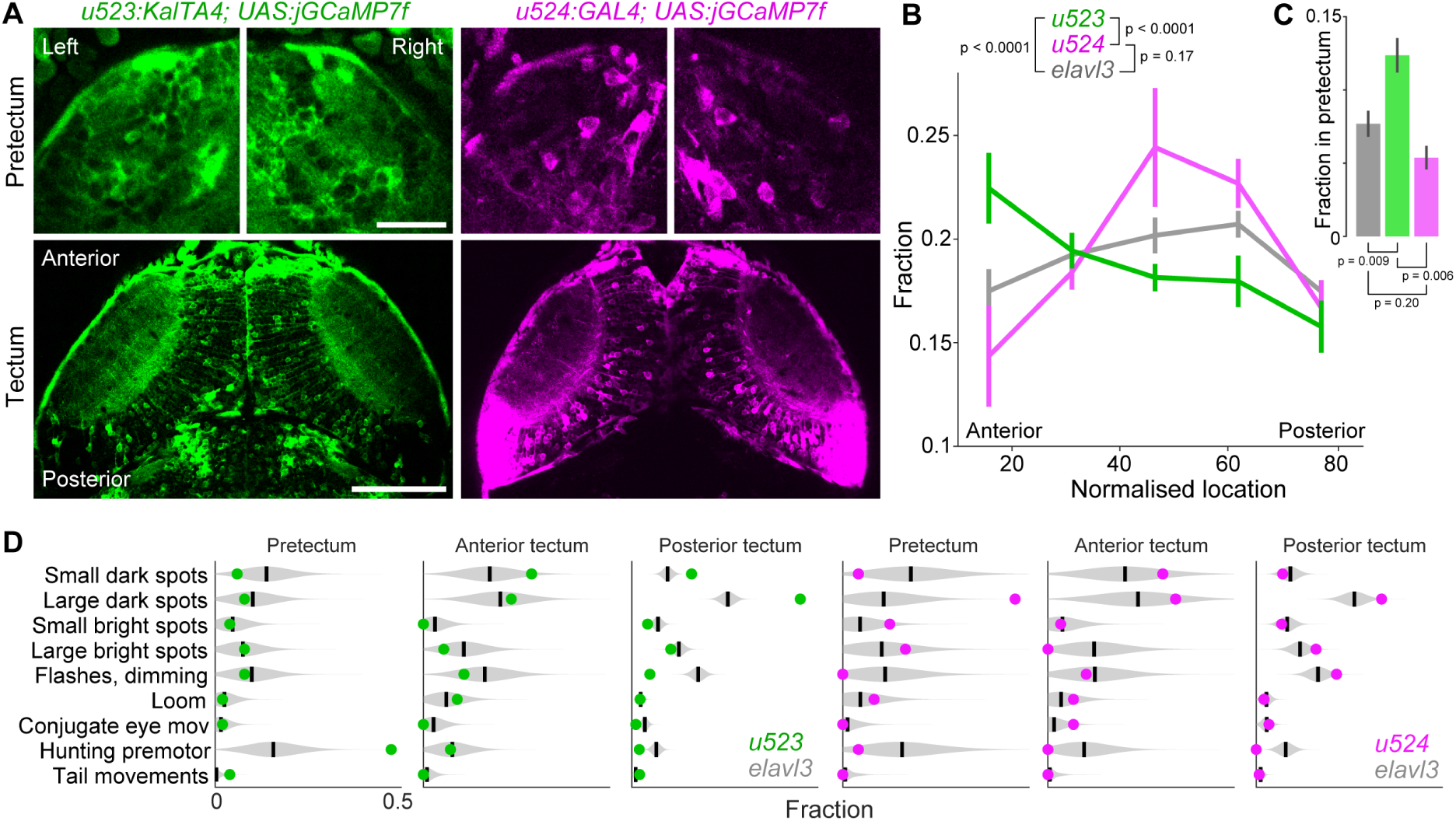
Transgenic lines labelling Pt7-OT neurons. (A) Optical sections of Pt7 (top) and OT (bottom) in a Tg(*u523*:KalTA4;UAS:jGCaMP7f) larva (green, left) and a Tg(*u524*:GAL4;UAS:jGCaMP7f) larva (magenta, right) at 6 dpf. Scale bars: Pretectum, 25 *µ*m; Tectum, 100 *µ*m. (B) Distribution of labelled cells across the anterior-posterior axis. Pt7 falls within the most anterior bin. Mean ± SEM was computed across fish (*N* = 11 *u523* larvae, *N* = 5 *u524*, *N* = 13 Tg(*elavl3*:H2B-GCaMP6s)). (C) Fraction of cells in Pt7 relative to total Pt7-OT cells in each line. *u523* preferentially labels pretectal neurons. (D) Functional properties of transgenically labelled neurons. Subplots show the fraction of cells assigned to each functional group for *u523*:GCaMP7f (left, green) and *u524*:GCaMP7f (right, pink) transgenics. Violin plots show reference distributions (lines indicate mean) generated by repeatedly (2000-fold) sampling, for each *u523* or *u524* cell, the cluster identity of one neuron from a group of 40 nearest-neighbour neurons (within 20 *µ*m in ZBB atlas space) labelled by Tg(*elavl3*:H2B-GCaMP6s).

Next, we used calcium imaging to assess the functional properties of transgenically labelled neurons. This revealed that *u523* and *u524* preferentially label particular ensembles of functional cell types, beyond what would be expected from the anatomical distribution of transgene expression alone. We determined this by computing the fraction of *u523* and *u524* cells belonging to different functional groups and compared these values to reference distributions generated by pseudorandom sampling of *elavl3*:H2B-GCaMP6s neurons at anatomically matched locations [Figure 3D; see Methods]. Of particular relevance to this study, the *u523* transgenic showed a substantial over-representation of hunting premotor neurons, with a rate of labelling in Pt7 that was almost four-fold greater than the *elavl3* reference distribution. In contrast, *u524* labeled very few of these neurons. Therefore, we next used the *u523* transgenic line for optogenetic control of hunting premotor activity.

### Optogenetic stimulation of Pt7 supports a rate code controlling the induction and steering of hunting behaviour

We used optogenetic stimulation to test the hypothesis that rate-coded activity in Pt7 controls the induction of hunting and steering of predatory actions. We expressed the excitatory opsin CoChR^32^ in pretectal neurons using either the *u523* driver line or the Tg(*u508*:KalTA4) (hereafter *u508*) line, which labels pretectal command neurons capable of inducing hunting^18^. We used a custom digital-micromirror-device (DMD) system to focally activate Pt7 in tethered larvae while tracking eye and tail movements using a high-speed camera [Figure 4A] and modulated stimulus irradiance as a means to evoke varying levels of pretectal activity.

**Figure 4:**
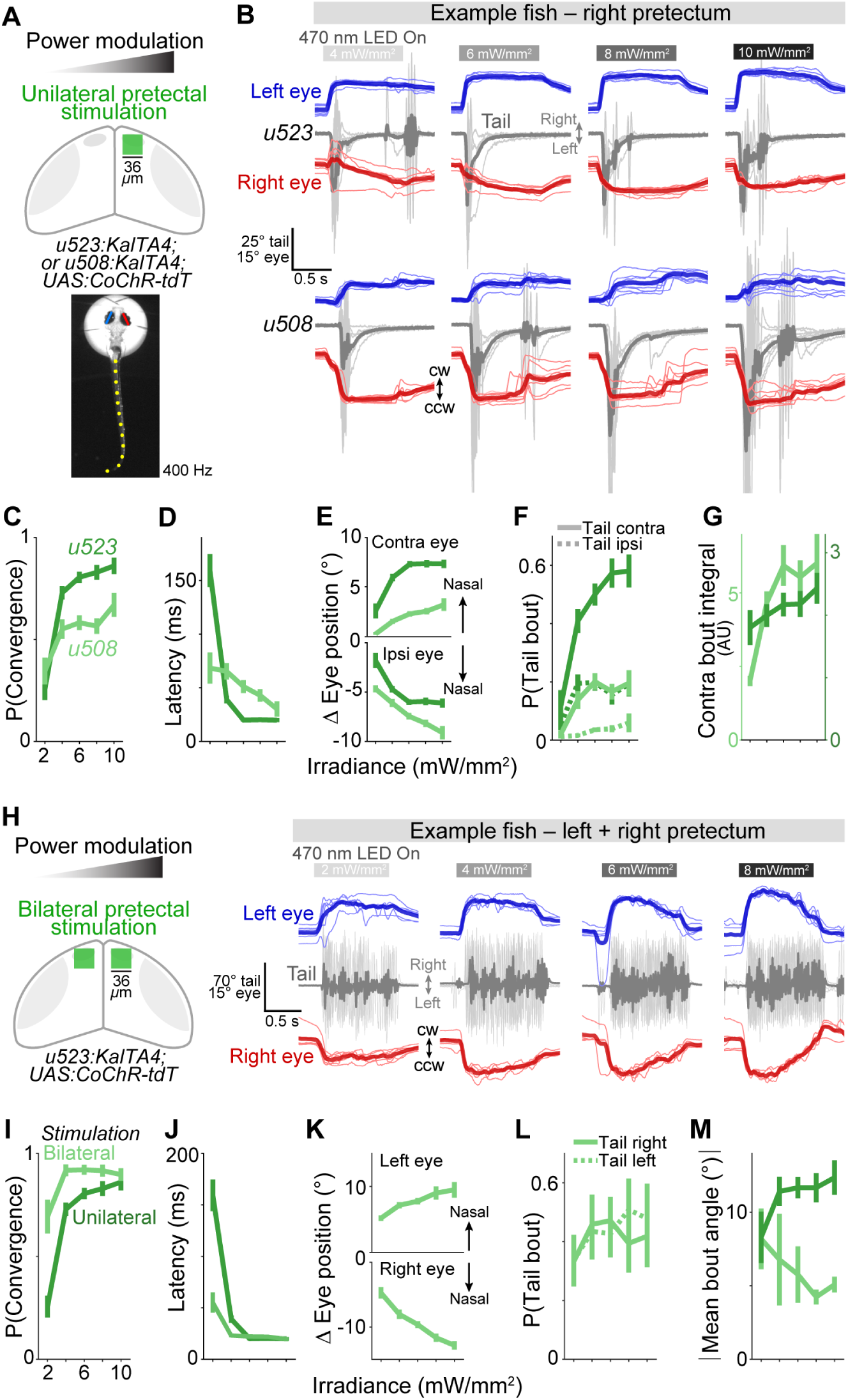
Optogenetic stimulation of Pt7 supports a rate code for control of hunting initiation and orientation. (A) Optogenetic stimulation. Pt7 was stimulated (470 nm; 36 × 36 *µ*m; 1 s LED-on) at varying irradiance (2 − 10 mW/mm^2^) and behaviour was tracked at 400 Hz. (B) Examples of optogenetically induced behaviour. Thin lines show individual trials, thick lines show mean. CW, clockwise; CCW, counter-clockwise. Stimulation of right Pt7 evokes a convergent saccade paired with a leftwards (negative) swim bout. (C) Probability of evoking a convergent saccade versus stimulus irradiance. Mean ± SEM was computed across fish, as for all other plots (*N* = 30 *u523* larvae, *N* = 47 *u508*). (D) Latency from stimulus onset to saccade onset. (E) Amplitude of evoked eye movements. Ipsi and contra relative to stimulated Pt7. (F) Probability of convergent saccade being paired with contraversive or ipsiversive tail movement. (G) Integrated tail bend angle during contraversive tail movements. (H) Bilateral Pt7 stimulation. (I-M) As per C-G but for bilateral stimulation, *N* = 9 larvae. Unilateral data overlaid for comparison.

Unilateral stimulation of Pt7 in both *u523* and *u508* larvae was effective at evoking hunting. Optogenetically evoked behaviour comprised a convergent saccade paired with a contralaterally directed swim bout [Figure 4B] and occurred with short (*<* 75 ms) latency [Figure 4D]. By contrast, control stimulations that were targeted outside the brain never induced convergent saccades and only rarely evoked conjugate or divergent eye movements that occurred at significantly longer latency, suggesting such responses were visually evoked [Figure S5]. In addition, Pt7 stimulation failed to induce hunting in *u524* larvae, in accordance with the infrequent labelling of hunting premotor neurons in this line [Figure S5].

In support of a rate code, increasing stimulation irradiance led to an increase in both the probability and steering of hunting actions. Specifically, the probability of evoking a convergent saccade increased monotonically from 2 to 10 mW/mm^2^ [Figure 4C]. Increasing stimulus intensity also resulted in a higher probability of a contralaterally directed tail movement [Figure 4F] and such movements displayed a progressive increase in integrated tail-bend angle [Figure 4G], indicative of a stronger orienting response.

To distinguish the influence of pretectal activity on the induction versus steering of hunting actions, we took advantage of the fact that forward-directed hunting responses are associated with symmetric activity across left and right Pt7-OT. Thus, we performed bilateral optogenetic simulation of Pt7 using the *u523* transgenic line [Figure 4H]. As before, increasing stimulus intensity led to an increase in the probability of evoking a convergent saccade [Figure 4I]. However, in contrast to unilateral stimulation, bilateral activation consistently generated symmetric tail movements resembling forward swims [Figure 4H]. Swim bouts were initiated towards left or right with equal probability [Figure 4L] and had low bend angle [Figure 4M], indicative of forward swimming with minimal lateralisation.

In sum, these results support a causal role for rate-coded activity in pretectal premotor neurons controlling both the induction of hunting and the steering of predatory actions. Increases in overall Pt7 activity increase the probability of initiating hunting and increasing lateralisation of Pt7 activity produces stronger contraversive orienting.

### Optogenetic stimulation of OT reveals pronounced anterior-posterior functional topography

Because calcium imaging revealed premotor activity in aOT and mOT contralateral to turn direction as well as activity in pOT on the ipsilateral side, we sought to test how this topographically patterned tectal activity shapes behaviour. To this end, we performed focal optogenetic stimulation at different sites along the anterior-posterior axis of OT in Tg(*u523*:KalTA4;UAS:CoChR-tdT) larvae [Figure 5A].

**Figure 5:**
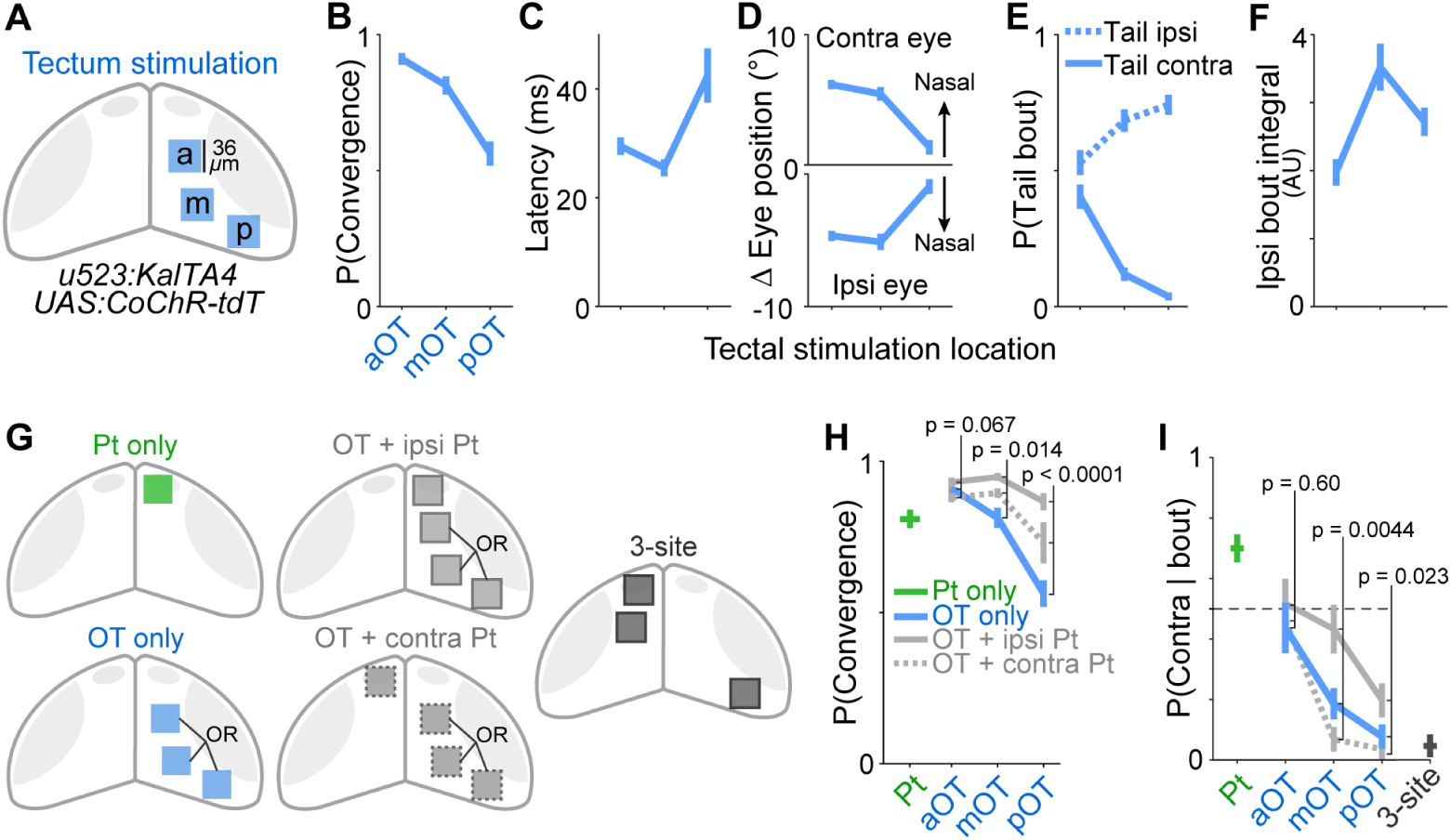
Optogenetic stimulation of Pt7 and OT supports a joint role in hunting control. (A) Optogenetic stimulation of anterior (a), middle (m), and posterior (p) tectal loci (470 nm; 8–10 mW/mm^2^; 36 × 36 *µ*m; 1 s LED-on). (B) Probability of evoking a convergent saccade versus tectal stimulation locus. Mean ± SEM was computed across fish, as for all other plots (*N* = 28 larvae). (C) Latency of saccade onset. (D) Amplitude of evoked eye movements. Ipsi and contra relative to tectal locus. (E) Probability of convergent saccade being paired with contraversive or ipsiversive tail movement. (F) Integrated tail bend angle during ipsiversive tail movements. (G) Schematic of combinatorial optogenetic stimulations. Loci in OT were stimulated in combination with either ipsilateral or contralateral Pt7. (H) Probability of evoking a convergent saccade. Data for stimulation of OT alone (blue) is reproduced from (B) and results of stimulating Pt7, alone (green) or in combination with OT (grey), are shown. (I) Probability that optogenetically induced tail movements were contralaterally directed. For co-activations, direction is relative to tectum.

The probability and direction of optogenetically evoked hunting outputs were acutely sensitive to the locus of tectal stimulation. The probability of evoking a convergent saccade was greatest in aOT and decreased significantly as stimulation was targeted to more posterior sites [Figure 5B]. As with Pt7 stimulations, optogenetically evoked convergent saccades were typically associated with tail movements. However, while aOT stimulations produced a similar frequency of contra- versus ipsiversively directed swims, more caudal loci evoked ipsiversive turns [Figure 5E]. Furthermore, the strength of ipsiversive swims was greatest following stimulation of m/pOT [Figure 5F]. As before, optogenetic stimulation using the *u524* driver line was not effective at evoking hunting [Figure S5].

In sum, these data support a model in which aOT contributes to the induction of hunting but imparts little directionality on predatory actions. By contrast, pOT is less effective at inducing hunting, but does have a strong directional influence, steering predatory swims ipsiversively.

### Combinatorial stimulation of Pt7 and OT supports a joint role in control of hunting

During naturalistic hunting, premotor neurons in both Pt7 and OT are recruited [Figure 1, Figure 2]. To investigate how activity in these regions interacts to command and pattern hunting responses, we stimulated Pt7 and OT simultaneously. Specifically, we activated loci at one of three locations along the anterior-posterior axis of OT in combination with either ipsi- or contralateral Pt7 [Figure 5G].

First, we assessed the induction of hunting. For mOT and pOT, additional stimulation of Pt7 significantly increased the likelihood of a hunting response [Figure 5H]. For aOT, response probability was already very high and there was no clear effect of additionally stimulating Pt7.

Next, we assessed the direction of tail movements following combinatorial stimulations. Additional stimulation of Pt7 had minimal effect when combined with aOT activation: As before, we observed a similar frequency of ipsi- versus contraversive tail movements [Figure 5I]. However, co-activation of Pt7 did influence steering when combined with mOT/pOT activation, with tectum and pretectum having opposite directional biases. Thus, Pt7 stimulation on the same side of the brain acted in opposition to mOT/pOT stimulation to reduce ipsilateral bend angle, whereas stimulating Pt7 in the opposite hemisphere augmented the effect of tectal stimulation to produce stronger ipsiversive (with respect to OT) turns [Figure 5I]. Consequently, the most strongly lateralised hunting responses were generated by stimulating pOT on one side of the brain and Pt7 on the opposite side. A similar result was produced by a three-site stimulation, designed to recapitulate the topography of premotor activity observed by calcium imaging. Specifically, we stimulated both Pt7 and aOT on one side as well as pOT in the opposite hemisphere and again observed strongly lateralised predatory orienting [Figure 5I, ‘3-site’].

In summary, focal optogenetic stimulation supports a causal role for topographically organised activity in Pt7-OT inducing and directing hunting behaviour. Next, we wanted to assess how the functions of different Pt7-OT sites was linked to their patterns of efferent projections to the hindbrain.

### Topography of descending projections aligns with the Pt7-OT motor map

Orienting and avoidance turns have long been associated with activity of crossed and uncrossed tectobulbar pathways^1^. The functional topography revealed by calcium imaging and optogenetics predicted that aOT might provide similar output into crossed and uncrossed descending pathways, while Pt7 should preferentially innervate contralateral hindbrain, and mOT/pOT is expected to show an ipsilateral bias.

To test these predictions, we examined axonal projections from circumscribed regions of Pt7-OT by performing multiphoton photoactivation of PA-GFP in Tg(*α*-tubulin:C3PA-GFP; *elavl3*:jRCaMP1a) double-transgenic animals at 5 dpf [Figure 6A,B]. Labelled neurites were segmented from imaging volumes acquired at 6–7 dpf and mapped to reference brain space using the jRCaMP1a channel for volumetric registration.

**Figure 6:**
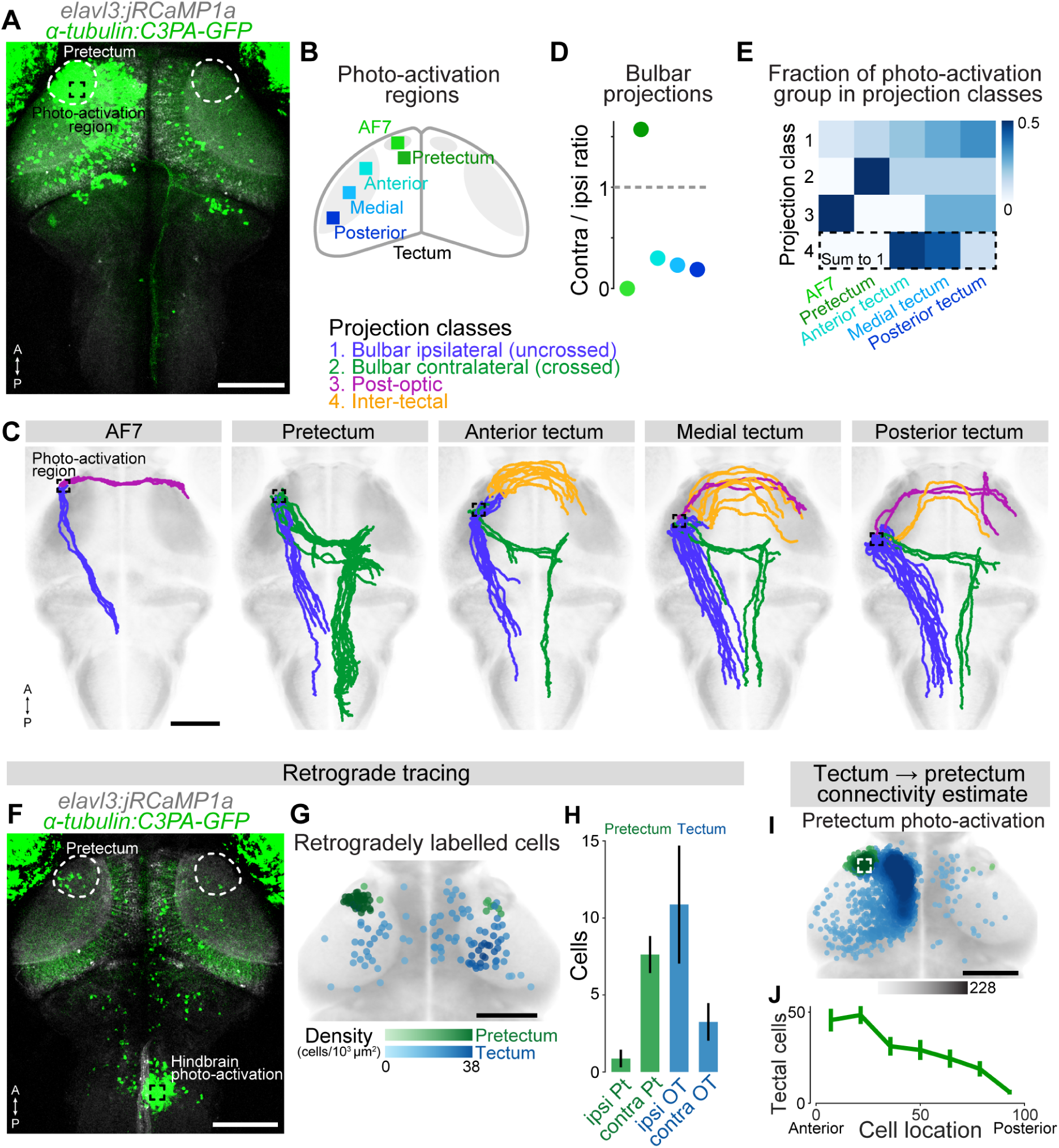
Topography of Pt7-OT projections. (A) Photoactivation of PA-GFP. Maximum intensity projection of Tg(*α*-tubulin:C3PA-GFP;*elavl3*:jRCaMP1a) larva at 6 dpf, following photoactivation in pretectum at 5 dpf. Photolabelled somata and neurites are visible in green, while jRCaMP1a expression is overlaid in grey. Scale bar: 100 *µ*m; A, anterior; P, posterior. (B) Schematic of photoactivation loci in Pt7-OT. At each site, a 9 × 9 × 40 *µ*m photo-activation volume was scanned at 790 nm. (C) Photolabelled neurites traced from each locus (black squares) and colour-coded according to projection class, *N* = 35 larvae. Scale bar: 100 *µ*m. (D) Ratio of contra-(class 2) to ipsilateral (class 1) bulbar projections deriving from each photoactivation locus. (E) Fraction of neurites belonging to each projection class that derive from each photoactivation locus. Values in each row sum to one. (F) Retrograde labelling of Pt7-OT somata. Maximum intensity projection at 7 dpf following photoactivation in ventral-medial hindbrain at 6 dpf. (G) Retrogradely labelled somata in pretectum (green) and tectum (blue) following photoactivation in hindbrain (*N* = 8 larvae). Darker colour indicates higher local cell density. Scale bar: 100 *µ*m. (H) Counts of retrogradely labelled cells. Mean ± SEM was computed across fish, as for other plots in this figure. (I) Retrogradely labelled somata in pretectum (green) and tectum (blue) following photoactivation in pretectum (*N* = 7 larvae). Scale bar: 100 *µ*m. (J) Distribution of retrogradely labelled tectal cells across the anterior-posterior axis following photoactivation in pretectum.

Using an unsupervised clustering procedure, we identified four major classes of projections from Pt7-OT [Figure S6A,B]: (1) Uncrossed bulbar pathway, with descending projections to ipsilateral ventral hindbrain; (2) Crossed bulbar pathway, with axons decussating near the oculomotor nucleus and extending into contralateral ventral-medial hindbrain; (3) Ventral inter-hemispheric pathway, projecting to contralateral Pt7-OT via the post-optic commissure; (4) Dorsal interhemispheric pathway, comprising bundles of axons projecting to the contralateral tectum via dorsal midbrain tracts. By comparing to previously described single-cell morphologies^6,18,19,33^ we verified that these classes correspond to mutually exclusive projection phenotypes of individual neurons [Figure S6C,D].

There was substantial topographic variation along the anterior-posterior axis of Pt7-OT in terms of efferent projections into the crossed and uncrossed bulbar pathways [Figure 6C-E]. To show this, we computed the ratio of class 2 to class 1 neurites from each photo-activation site, which revealed that pretectum primarily makes crossed descending projections, whereas most axons originating from OT enter the uncrossed tectobulbar pathway [Figure 6D]. Additionally, we estimated the fractional contribution of each photoactivation region to each pathway [Figure 6E], which indicated that pretectum is the largest contributor to the crossed bulbar pathway (55% of class 2 neurites) whereas an increasingly large proportion of uncrossed bulbar projections derive from more caudal loci in OT (20, 26 and 32% from a-, m- and pOT respectively). Because photoactivation is likely to label axons of passage as well as projections originating from cells in the photoactivation volume, we sought to corroborate these findings by performing retrograde labelling. Accordingly, we performed unilateral photoactivations in a region of the ventromedial hindbrain neuropil where both crossed and uncrossed bulbar axons extend [Figure 6F] and counted retrogradely labelled cell bodies in Pt7-OT on both sides of the brain [Figure 6G]. This confirmed that pretectal neurons primarily project contralaterally whereas most descending projections from OT are uncrossed/ipsilateral [Figure 6H].

### Reciprocal connectivity between Pt7 and OT

Previous findings suggest strong interconnections between aOT and Pt7. First, optogenetic stimulation of anterior-ventral OT can induce hunting actions^21^, but this is abrogated by ablation of pretectal command neurons^18^, suggesting aOT recruits Pt7 to induce hunting. Second, in the opposite direction, a subset of pretectal command neurons make ipsilateral projections into aOT^18^.

To further explore reciprocal connectivity, we identified retrogradely labelled cell bodies following photoactivation at each Pt7-OT locus [Figure S6E-G]. This revealed a large number of cells in ipsilateral aOT that extend neurites into Pt7, but many fewer such cells at more caudal tectal locations [Figure 6I,J]. In support of reciprocal connectivity, ipsilateral pretectal neurons were retrogradely labelled from aOT and, to a lesser extent, from mOT [Figure S6F].

In sum, these anatomical data reveal pronounced topography in Pt7-OT projection patterns, which align with the motor map described above. Pt7 makes a conspicuous projection to the contralateral hindbrain, in line with our finding that premotor neurons in this region evoke contraversive predatory orienting. Strong reciprocal connections with aOT provide a putative anatomical substrate for the recruitment of premotor activity across Pt7-aOT during hunting behaviour [Figure 2] as well as the observation that additional co-activation of Pt7 does not augment hunting induction beyond stimulating aOT alone [Figure 5]. In OT, we observed an anterior-posterior gradient in descending connections wherein more posterior loci increasingly project to ipsilateral hindbrain in agreement with the fact that these loci evoke the strongest ipsiversive hunting manoeuvres.

### Pretectum controls contraversive orienting via the crossed pretectobulbar pathway

Finally, we sought to verify that Pt7 projections to the contralateral hindbrain do in fact underlie contraversive hunting manoeuvres. To do this, we combined optogenetic stimulations with laser-ablation of crossing (type 2) pretectobulbar axons. We first photoactivated PA-GFP bi-laterally in Pt7 in triple-transgenic Tg(*u523*:KalTA4; UAS:CoChR-tdTomato; *α*-tubulin:C3PA-GFP) larvae at 5 dpf and subsequently laser-ablated photolabelled axons crossing the midline at 6 dpf [Figure 7A-C]. For each animal, we performed optogenetic stimulation and behavioural tracking both immediately prior to axotomy and following overnight recovery.

**Figure 7:**
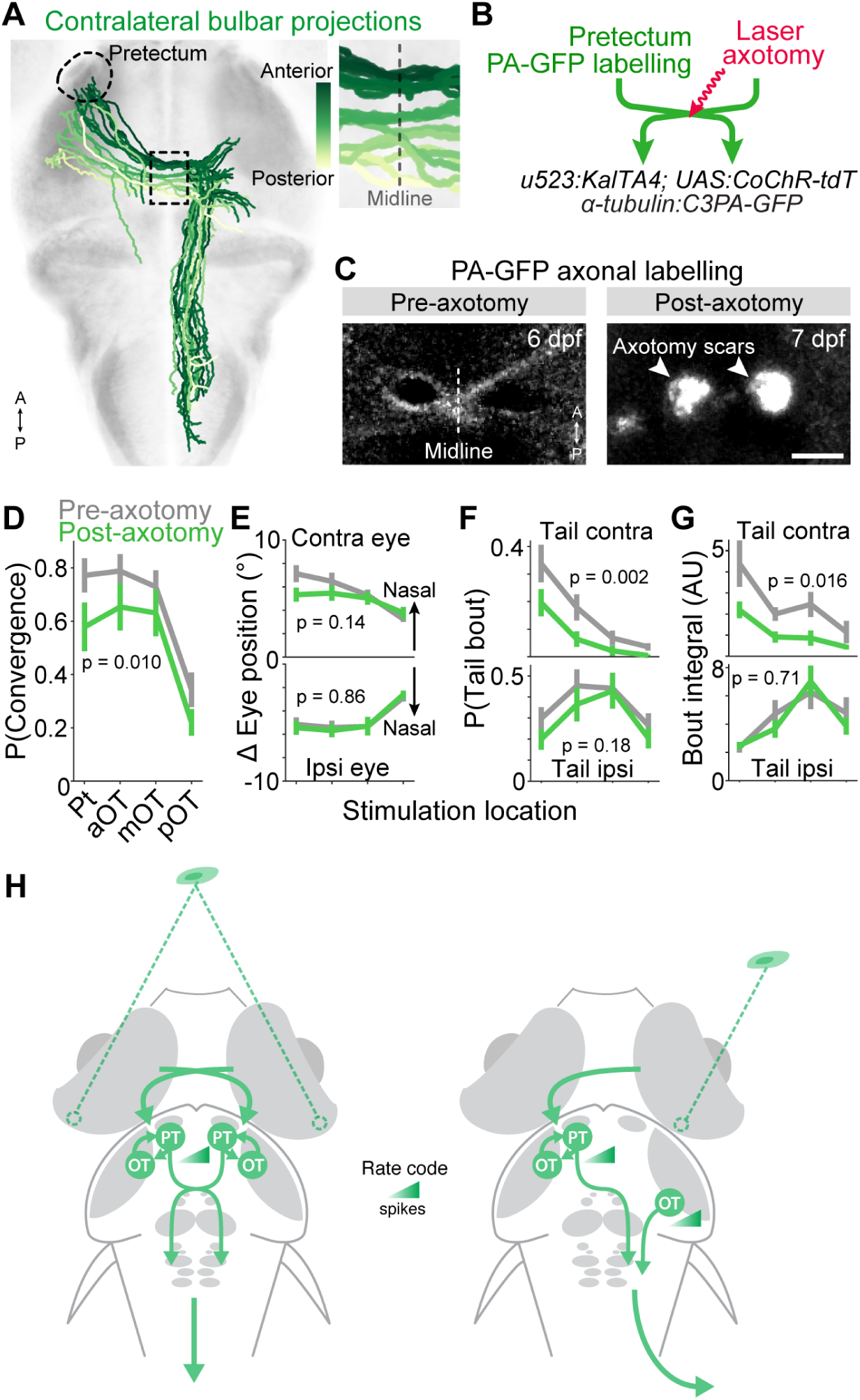
Crossed pretectobulbar projections mediate contraversive hunting actions. (A) Crossed projections (class 2) colour-coded according to anterior-posterior location of midline decussation (enlarged in inset). Axons deriving from pretectum cross at more anterior locations. (B) Schematic of laser axotomies. (C) Maximum intensity projections of midline region showing PA-GFP-labelled pretectobulbar axons prior to (left, 6 dpf) and after (right, 7 dpf) axotomy. The autofluorescent ‘scars’ typical of multiphoton ablation are clearly visible. Scale bar: 15 *µ*m. (D) Probability of optogenetically evoking a convergent saccade versus stimulus location before (grey) and after (green) laser axotomy. Mean ± SEM was computed across fish, as for all other plots (*N* = 18 larvae). (E) Amplitude of evoked eye movements. Ipsi and contra relative to stimulation site. (F) Probability of convergent saccade being paired with contraversive or ipsiversive swim. (G) Integrated tail bend angle for contraversive and ipsiversive tail movements. In D-G, *p*-values from two-way ANOVA comparing pre- vs post-axotomy. (H) Model for Pt7-OT control of predatory orienting. See main text for explanation.

Ablation of pretectobulbar axons reduced both the induction and contralateral steering of hunting actions. Specifically, the probability of optogenetic stimulation inducing a convergent saccade was reduced following axotomy [Figure 7D], with the greatest effect for stimulation of anterior sites (Pt7 and aOT). Moreover, we observed a decrease in the frequency and amplitude of contraversive tail movements coincident with convergent saccades, whereas ipsiversive tail movements were not significantly affected [Figure 7F-G]. We also attempted to lesion the uncrossed tectobulbar pathway but did not observe changes in optogenetically evoked behaviour, which we suspect is due to the difficulty of ablating this broad and dense axon tract [Figure S7].

In sum, loss-of-function experiments are consistent with crossed pretectobulbar projections mediating contraversive predatory orienting.

### A model for premotor control of predatory orienting

Together, our findings support a model in which rate- and space-coded premotor activity in Pt7 and OT jointly controls predatory orienting [Figure 7H]. In this model, hunting state is induced by activation of Pt7 command neurons^18^, which receive direct retinal input (to AF7) as well as input from aOT via strong reciprocal connections. While symmetric Pt7-aOT activity evokes forward-directed hunting manoeuvrers to centrally located prey targets, lateralised activity generates contraversive predatory orienting towards eccentric targets as a result of a predominantly crossed descending projection to contralateral hindbrain. Increased premotor activity generates higher amplitude orienting movements. In addition, activity in pOT in the opposite hemisphere augments the orienting response, acting via uncrossed tectobulbar projections.

## Discussion

Here, we present evidence that space- and rate-coded activity of premotor neurons in Pt7 and OT controls the steering of target-directed actions during hunting. Our key findings are consistent with: i) pretectum and OT jointly controlling the induction and steering of hunting behaviour; ii) both space- and rate-coded premotor activity controlling the amplitude of eye and body movements; iii) both crossed and uncrossed (pre)-tectobulbar pathways contributing to predatory orienting. Our discussion will focus on these aspects.

### Pretectum and optic tectum jointly control the induction of hunting state and predatory orienting

Together with earlier studies, our data indicate a close functional coupling between Pt7 and OT in both the induction of the hunting programme and the directional control of predatory actions. While the involvement of SC/OT in predatory orienting is well established^1^, a key finding of the present study is that Pt7 also contributes to direction control of eye and body movements towards prey targets. Our previous work established a crucial role for Pt7 in the induction of hunting state^18^. A subset of Pt7 cells labelled by the *u508* transgene are specifically active in association with hunting, their optogenetic activation can induce complete hunting sequences (in the absence of prey), and their ablation impairs natural hunting. Thus, these cells function as a forebrain command system controlling of engagement of hunting behaviour. Hunting can also be evoked by stimulation of aOT^21^, but this requires an intact Pt7 command system^18^, indicating that tectal activity sits upstream of Pt7. The extensive reciprocal connectivity we identified between aOT and Pt7 provides the likely circuit substrate for these interactions.

If Pt7-OT premotor activity controls both the induction and steering of hunting behaviour, how can they be differentially controlled, for example to respond with high probability to centrally located targets? At first glance, the strong Pt7-aOT activity that underlies a high response probability would also generate an inappropriate, high-amplitude orientation. The solution appears to come from a bilateral pattern of activity, observed by calcium imaging and shown with optogenetics to produce forward-directed hunting responses. Such activity likely provides ‘balanced’ premotor drive to tegmental circuits resulting in a less lateralised response as compared to that produced by one side alone.

### Space and rate-coded Pt7-OT activity controls target-directed orienting

Seminal studies in monkeys and cats have given rise to a model of collicular function in which the location of a ‘bump’ of premotor activity is the key variable that specifies a certain gaze displacement vector, while the level of activity influences the velocity (but not amplitude) of gaze shifts^34–41^. Because this motor map is aligned to topographically organised sensory representations, this arrangement has been assumed to enable a simple sensorimotor transformation across the depth of SC/OT to direct gaze towards targets of interest^1,2^ (but see also^42^). Evidence for such a space code has been found in a broad variety of vertebrate species ^1^. While individual premotor neurons have broad movement fields, the gaze displacement vector can nonetheless be accurately decoded from the population of active neurons^2,36,38^. In agreement with this longstanding model, we previously showed that predatory convergent saccades are associated with spatially localised bursts of activity in OT (tectal ‘assemblies’), whose anterior-posterior location correlates with saccade amplitude^14^. Our present results extend these findings, revealing a motor map across Pt7-OT in which the centroid of a broad bump of premotor activity covaries with the amplitude of both eye and tail movements.

Orienting movements are generated by rate-coded activity in motoneurons and it is assumed that the necessary transformation of space-coded activity in SC/OT to rate-coded activity occurs in the tegmentum^2^. Anatomical tracing in cats is consistent with this spatio-temporal transformation being achieved by a gradient of synaptic connection strengths^43^, while our data provides evidence that the *number* of tectal neurons projecting to the tegmentum also shows a spatial gradient along the anterior-posterior axis of Pt7-OT. However, our findings also indicate that rate-coded premotor activity controls the amplitude of orienting movements already at the level of Pt7-OT. This aligns with previous evidence from several species, in particular in relation to orienting movements of the head/body (rather than the eyes). For example, electrical microstimulation of SC/OT in goldfish ^44^, rodents^3,45^, and owls ^46^ has demonstrated that stimulation intensity, as well as location, influences the amplitude of head/body movements. Our data suggests that hunting premotor neurons in Pt7-OT linearly encode the amplitude of eye and body displacements. Although calcium imaging has limited sensitivity and we analysed inferred spike counts that are unlikely to accurately reflect firing rates, our results align with electrophysiological recordings from cats^38^, where tectoreticular neurons also display ‘open’ movement fields in which activity increases monotonically with movement amplitude in the cells’ preferred direction. Further support comes from recent recordings in mice, where the firing rates of tectoreticular neurons encode intended head displacements^47^. The fact that (pre)tectal activity might encode orienting movements using a rate code does not, however, guarantee that the decoding of premotor activity shows any dependency of activity level. Here, our optogenetic stimulation results are key, as we show that stimulation of the same locus, while only increasing irradiance, produces smooth, monotonic increases in the amplitude of eye and tail movements. Importantly, we used long stimulations that greatly outlasted the duration of behavioural responses. This strategy ensured that under conditions of space-coding, the extended stimulation would be sure to evoke a plateau response having the site-specific characteristic vector^39^. Small movements, evoked at low irradiance, did not increase in amplitude over the course of extended illumination, implying a true dependency on the level of evoked activity. Taken together, our imaging and optogenetic results support a role for both space- and rate-coded premotor activity in controlling the amplitude of eye and tail movements during predatory orienting.

### A joint role for crossed and uncrossed (pre)-tectobulbar pathways

Of the two major descending pathways from OT, there is longstanding evidence from a variety of species that cTB supports contraversive orienting, whilst iTB is associated with ipsiversively directed escape/avoidance^1,2^. However, two lines of evidence add complexity to this simple picture. First, animals can produce different types of orienting behaviour, which appear to have different dependencies on crossed and uncrossed descending pathways^3,48–50^. Dean et al.^3^ distinguished hunting-related ‘pursuit’, which enables animals to orient towards unpredictably moving targets under closed-loop control, from ‘localisation’-type movements in which orienting is guided to the remembered location of a transient stimulus in the periphery in an open-loop manner. In agreement with our model, lesion studies in rodents, cats, and frogs provide evidence that hunting-related pursuit is dependent on an intact cTB pathway, whilst localisation-type orienting can be sustained by alternative tectofugal circuits^3,51–53^. Second, although cTB might be required for normal predatory orienting, some studies suggest at least some role for iTB as well ^5,6,54,55^ . For example, multi-site lesions in frogs ^5^ provided evidence that uncrossed tectofugal pathways can generate predatory orienting and in agreement with our study, movements were ipsiversively directed (relative to OT). However, the responses were misdirected with respect to the visual target and it was unclear if/when the uncrossed pathway was activated during normal behaviour. More recently, optogenetic stimulation of a neuropil region containing iTB fibres was shown to evoke small contraversive tail movements in zebrafish^6^. Although opposite in directionality as compared to our model of iTB function, the relevance of these movements to hunting was unclear and so they might represent a parallel pathway or perhaps contribute to a different class of orienting behaviour.

By using convergent saccades as a robust indicator of the engagement of hunting state^9,10,14,23^ alongside transgenic lines enriched for hunting premotor neurons, we specifically interrogated the directional control of predatory orienting. Broad-scale calcium imaging across both tectal hemispheres during naturalistic behaviour, optogenetic stimulation, and projection tracing collectively support a model in which both cTB (originating primarily from Pt7) and iTB (from pOT in the opposite tectal hemisphere) cooperate to steer predatory actions. While the two tectofugal pathways generate predatory orienting with opposite directional biases with respect to OT, their recruitment in opposite hemispheres produces a cooperative action to direct eye and body movements towards prey targets. A joint role for crossed and uncrossed pathways also aligns with recent viral tracing studies in mammals which show that individual tectal projection neurons belonging to the ‘orienting’ pathway in fact extent bilateral descending axon collaterals^56^. Although our proposed roles for cTB and iTB are supported by spatially patterned optogenetic stimulation alongside the topography of descending projections, we did not specifically activate neurons contributing to defined tectofugal pathways. In future studies, our model should therefore be tested using projection-specific stimulation^56^.

### Bi-hemispheric premotor control

Our data show that predatory orienting — towards targets at either central or peripheral locations — involves activation of multiple loci spanning both brain hemispheres. Forward-directed responses associated with symmetric premotor activity in Pt7-aOT, which likely provides ‘balanced’ premotor drive to tegmental circuits to generate forward swims. This premotor activity lies in close proximity to visually responsive neurons with binocular receptive fields ^57^ and interactions between these two sets of neurons might enable depth-dependent control of prey pursuit^11^. The interhemispheric projections we traced between left and right Pt7-aOT might provide a circuit basis for binocular integration and/or balancing of premotor drive.

For large amplitude orientations to eccentric prey, an unexpected finding was recruitment of strong premotor activity in pOT, ipsilateral to turn direction. Because the retinotectal projection is entirely crossed in larval zebrafish, this ipsilateral activity must result from interhemispheric connections. Moreover, the coincidence of pOT activity with much more rostral activity in the opposite hemisphere suggests that any such connections do not simply link tectal loci with mirror-symmetric retinotopic coordinates. Possible pathways include direct projections between left and right OT^6^ or more indirect routes, for example via the nucleus isthmi^25^ or intertectal neurons^58^ and will be an important subject of future investigation.

## Acknowledgements

The authors thank lab members for helpful discussions and critical feedback on the project, and the UCL Fish Facility staff for fish care and husbandry. This research was funded in whole, or in part, by the Wellcome Trust (grant numbers 101195/Z/13/Z and 220273/Z/20/Z awarded to I.H.B.). For the purpose of Open Access, the author has applied a CC BY public copyright licence to any Author Accepted Manuscript version arising from this submission.

## Author Contributions

Conceptualisation: P.A. and I.H.B.; Investigation: P.A. and S.C-F.; Formal Analysis: P.A.; Writing: P.A. and I.H.B.; Supervision and Funding Acquisition: I.H.B.

## Methods

### Zebrafish lines and care

Zebrafish lines were maintained in the Tübingen background. Larvae were reared in fish-facility water on a 14/10 h light/dark cycle at 28.5*^◦^*C and were fed *Paramecia* from 4 dpf onwards. All larvae were homozygous for the *mitfa^w^*^2 59^ skin-pigmentation mutation. For calcium imaging, animals were either Tg(*elavl3*:H2B-GCaMP6s)jf5Tg ^60^ or double-transgenic for Tg(UAS:jGCaMP7f)u341Tg ^18^ and either Tg(*-2.5pvalb6*:KalTA4)u523Tg ^61^ or Tg(*-2.4efnB2a*:Gal4FF)u524Tg (generated in this study, see below). Larvae used for optogenetic stimulation were double-transgenic for Tg(UAS:CoChR-tdTomato)u332Tg ^32^ and either Tg(*pvalb6*:KalTA4)u523Tg, Tg(*-2.4efnB2a*:Gal4FF)u524Tg, or Tg(*pvalb6*:KalTA4)u508Tg ^18^. Photoactivations were performed using larvae carrying Tg(*Cau.Tuba1*:c3PA-GFP)a7437Tg ^62^ and Tg(*elavl3*:jRCaMP1a)jf16Tg ^63^. Larvae used for laser axotomies were triple-transgenic for Tg(*Cau.Tuba1*:c3PA-GFP)a7437Tg, Tg(*pvalb6*:KalTA4)u523Tg, and Tg(UAS:CoChR-tdTomato)u332Tg. The sex of the larvae is not defined at the early stages of development used for these studies. Experimental procedures were approved by the UCL Animal Welfare Ethical Review Body and the UK Home Office under the Animals (Scientific Procedures) Act 1986.

### Generation of transgenic zebrafish

The Tg(*-2.4efnB2a*:Gal4FF)u524Tg (i.e. *u524*) line was isolated as follows. First, we used Gateway cloning (Invitrogen) to construct an expression vector in which ∼ 2.4 kb of zebrafish genomic sequence upstream of the start codon of the ephrin-B2a gene was placed upstream of the Gal4FF open reading frame. The genomic sequence was cloned using the following primers and Phusion PCR polymerase (Thermo Fisher Scientific):

- GGGGACAAGTTTGTACAAAAAAGCAGGCTtcccgttgggatgacttcttgtg
- GGGGACCACTTTGTACAAGAAAGCTGGGTaaaggtcttgtggagtgctggctgtg

(where capital letters indicate the attB1/B2 extension sequences). The expression vector was then microinjected into one-cell stage Tg(UAS-E1b:Kaede)s1999t ^64^ embryos at 30 ng/ml along with tol2 mRNA (30 ng/ml) and adult fish were screened for germline transmission by outcrossing and examining embryos for Kaede fluorescence. Positive embryos from a single fish were then raised to adulthood. This expression vector generated a range of expression patterns, one of which was designated the allele u524Tg.

### Two-photon calcium imaging and behavioural tracking

Larvae were tethered in 3% low-melting point agarose gel in a 35 mm petri dish lid and sections of gel were carefully removed using an opthalmic scalpel to allow free movement of the eyes and tail below the swim bladder. Larvae were allowed to recover overnight before testing at 6 or 7 dpf. Imaging was performed using a custom-built multiphoton microscope [Olympus XLUMPLFLN ×20 1.0 NA objective, 580 nm PMT dichroic, bandpass filters: 510/84 (green), 641/75 (red) (Semrock), R10699 PMT (Hamamatsu), Chameleon II ultrafast laser (Coherent)] at 920 nm with laser power at sample of 5–10 mW. Images (0.61 *µ*m/px) were acquired by frame scanning at 3.6 Hz, with focal planes separated by 8 − 10*µ*m.

Visual stimuli were back-projected (Optoma ML750ST) onto a curved screen placed in front of the animal at a viewing distance of 25 mm while a second projector provided constant background illumination below the fish. Visual stimuli were defined using the ‘red’ colour channel and Wratten filters (Kodak, no. 29) were placed in front of both projectors to block residual light that might be detected by the PMT. Visual stimuli were designed in MATLAB using Psychophysics Toolbox ^65^ and presented on a uniform grey background. Prey-like moving spots comprised 4, 8 or 16*^◦^* bright or dark spots (Weber contrast +1 or -1 respectively) moving at 30 or 80*^◦^*/s either left→right or right→left across 152*^◦^* of frontal visual space. Stimulus design took account of distortion caused by the curved screen to ensure constant angular velocity and angular size from the point of view of the fish. Looming stimuli comprised expanding dark spots (Weber contrast -1) that simulated an object approaching at constant velocity and were presented in front of the fish (10*^◦^*–70*^◦^*, L/V 490 ms) ^66,67^. We also presented a control dimming stimulus that had a fixed angular size equal to the final size of the looming spot (70*^◦^*) and dimmed so as to produce an identical change in whole-field luminance as the expanding looming spots. Finally, 3 s whole-field bright/dark flashes were presented. For all experiments, stimuli were presented in a pseudo-random sequence with 30 s inter-stimulus interval during which a uniform grey screen was shown.

Eye position was monitored at 60 Hz using a FL3-U3-13Y3M-C camera (Point Grey) that imaged through the microscope objective under 720 nm illumination. Tail position was imaged at 430 Hz under 850 nm illumination using a sub-stage GS3-U3-41C6NIR-C camera (Point Grey). Horizontal eye position and tail posture (defined by 13 equidistant x-y coordinates along the anterior-posterior axis) were extracted online using machine vision algorithms ^14^. Microscope control, stimulus presentation and behaviour tracking were implemented using LabView (National Instruments) and MATLAB (MathWorks).

### Analysis of behavioural data

Eye tracking data was processed to detect and classify saccadic eye movements as per ^23^. Hunting probability was computed as the fraction of trials in which at least one convergent saccade was detected during visual stimulus presentation or optogenetic stimulation. Response latency was calculated from stimulus onset. Post-saccadic version was computed as the mean across left and right eyes of their median position during a 200 ms interval following saccade detection.

Tail kinematics and swim bout detection were performed as per ^68^. In brief, 13 x-y centroids defined the midline of the tail, such that consecutive centroids defined tail ‘segments’. Vectors of 11 inter-segment angles were computed for each time-point and cumulative tail bend angle was computed as the sum of these inter-segment angles. Rightward bending of the tail is represented by positive angles and leftward bending by negative angles. Swim bouts were detected from the time-series of cumulative tail bend angle ^68^. Bout tail angle is the mean value of cumulative tail bend angle during the first 120 ms from bout onset. Bout integral is the summed value of cumulative tail bend angle for the full duration of the swim bout (divided by 1,000 for plotting purposes).

### Analysis of calcium imaging data

Motion correction of fluorescence imaging data was performed as per ^14^. Regions of interest (ROIs) corresponding to cell nuclei and somata were identified using either the cell detection algorithm of Kawashima et al. ^69^ for Tg(*elavl3*:H2B-GCaMP6s) data or Suite2P ^70^ for Tg(*u523*:KalTA4; UAS:jGCaMP7f) and Tg(*u524*:Gal4FF; UAS:jGCaMP7f) datasets. Time-varying fluorescence, *zF*, was computed as the mean value of all pixels within the ROI mask at each time-point (imaging frame) and normalised by z-scoring.

We used linear regression to model *zF* for each neuron. We designed 110 regressors, derived from behavioural and stimulus predictors [Table S1]. Visual stimulus predictors were binary vectors indicating the frames during which specific stimuli were presented. For each type of moving spot stimulus, we generated four predictors, each representing a quarter of azimuth space. Oculomotor predictors included left/right ipsiversive eye position, leftward/rightward conjugate saccades, and leftward/rightward convergent saccades. For the eye position predictors, nasal eye positions (estimated as those positions more nasal than the median position across the entire experiment) were zeroed. The saccadic predictors were one-hot encodings indicating the imaging frames corresponding to saccade onset. Locomotor predictors included leftward/rightward tail movements and tail vigour (smoothed absolute value of time derivative of cumulative tail bend angle). To account for indicator dynamics, regressors were generated by convolving each predictor with a calcium impulse response function (CIRF) modelled as an exponential rise and subsequent decay with time constants *τ_on_* = 20 ms and *τ_off_* = 2 or 3 s for jGCaMP7f or H2B-GCaMP6s, respectively. In addition, to capture fluorescence modulations that might result from residual motion artefacts, we included a ‘motion-error’ regressor, derived from the translation applied during motion correction of image time series (this regressor was not convolved with the CIRF). For each cell, we time-shifted the regressor matrix relative to the *zF* response variable so as to minimise the residual squared error of an ordinary least squares regression model. Then, we fit a linear regression model using elastic-net regularised regression ^71^. In practice, we used the MATLAB implementation of glmnet ^72^ with hyperparameters selected by 10-fold cross-validation. Fit model coefficients for oculomotor, locomotor, and motion error regressors (but not visual stimulus regressors) were used to construct visuomotor vectors.

Visuomotor vectors (VMVs) ^18^ were generated for each neuron by concatenating the following: (a) The integral of *zF* in response to each visual stimulus type. The median value across stimulus presentations was used and we only considered ‘no-response’ presentations in which the larva did not generate a convergent saccade (components 1–28). (b) Fit regression model coefficients (*β*s) for oculomotor, locomotor, and motion error regressors (components 29–38). VMVs from all neurons (83,656 cells from 29 larvae) were concatenated into a matrix and each component was normalised across cells by dividing by its standard deviation. To find groups of neurons with similar visuomotor responses, we performed hierarchical agglomerative clustering using a correlation distance metric ^14^. VMVs were embedded in a 2D space by applying t-distributed stochastic neighbour embedding (tSNE) using the Barnes-hut algorithm, a correlation distance metric and the following hyperparameters: perplexity 230, exaggeration 6, learning rate 500, theta 0.5.

We generated control distributions for functionally defined cell types as follows. For each *u523* or *u524* cell, we assessed the cluster identity of a randomly selected neuron from a group of 40 nearest-neighbours within 20*µ*m in ZBB space labelled by Tg(*elavl3*:H2B-GCaMP6s), and repeated this process 2000 times.

A Hunting Index (HIx) ^18^ was computed for individual neurons as a means to capture activity modulation specifically associated with the initiation of hunting behaviour (see schematic in [Figure S4]). For each stimulus-evoked convergent saccade, we first compute *x_Ri_* as the integral of *zF* during a time window (-1 to +2 s) relative to saccade onset time. Next, the integral of *zF* was extracted for the same time interval (with respect to stimulus presentation) during non-response trials in which the same visual stimulus was presented. The difference between *x_Ri_* and the mean of non-response activity (*µ_NR_*) was measured in units of the standard deviation of the non-response distribution (*σ_NR_*):

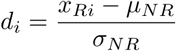

HIx scores were computed as the median of these *d_i_* distance values across all hunting responses during which the neuron was imaged and were computed separately for convergent saccades having leftward or rightward post-saccadic gaze.

To account for indicator dynamics and estimate ‘spiking’ activity, *zF* was deconvolved using OASIS ^28^ with a first-order autoregressive model. Convergent saccade-triggered activity was taken as the sum of OASIS-inferred spikes during a time window (±0.8 s) spanning saccade onset time.

Receptive field centres of spot-selective neurons (clusters 1–31) were estimated as follows. Spiking was assessed during presentations of moving spot stimuli for which the normalised VMV component exceeded one standard deviation. For time-points having a non-zero number of spikes, we determined the azimuth location of the spot and these azimuth values were gaze-adjusted by subtracting the position of the eye opposite to the brain hemisphere in which the cell was located (i.e. left eye for right-sided neurons).

The receptive field centre was estimated as the median of gaze-adjusted azimuth values weighted by the number of inferred spikes and receptive field width was computed as twice the median absolute deviation of this distribution. The estimated receptive fields of Pt7-OT neurons (34.3±11.8*^◦^*, mean ±SD, *N* = 5466 cells) were comparable with previous reports ^16,20,73^.

The centre of mass of tectal population activity was computed as the median normalised anterior-posterior location across neurons, weighted by each cell’s mean number of spikes for a given amplitude bin.

### Image registration

Registration of image volumes was performed using the ANTs toolbox version 2.1.0^74^ in a similar manner to that described in ^25^. Images were converted to NRRD file format for registration using ImageJ. As an example, to register the 3D image volume ‘fish1.nrrd’ to reference brain ‘ref.nrrd’, the following command was used:

~~~
antsRegistration -d 3 --float 1 -o [fish1,fish1_Warped.nii.gz] -n BSpline -r
[ref.nrrd,fish1.nrrd,1] -t Rigid[0.1] -m GC[ref.nrrd,fish1.nrrd,1,32,Regular,0.25]
-c [500×250×200×100,1e-8,5] -f 12×8×4×2 -s 4×3×2×1 -t Affine[0.1] -m
GC[ref.nrrd,fish1.nrrd,1,32,Regular,0.25] -c [500×250×200×100,1e-8,10] -f 12×8×4×2
-s 4×3×2×1 -t SyN[0.1,6,0] -m CC[ref.nrrd,fish1.nrrd,1,2] -c [200×200×200×200,1e-7,10]
-f 12×8×4×2 -s 4×3×2×1
~~~

The deformation matrices computed above were then applied to any other image channel N of fish1 using:

~~~
antsApplyTransforms -d 3 -v 0 -float -n BSpline -i fish1-0N.nrrd -r ref.nrrd -o
fish1-0N_Warped.nii.gz -t fish1_1Warp.nii.gz -t fish1_0GenericAffine.mat
~~~

Image volumes were registered to the Zebrafish Brain Browser (ZBB) atlas ^26,27^ and/or the Mapzebrain atlas^33^ with some differences between experiments:

- For calcium imaging volumes, a multi-step registration procedure was used. The functional imaging volume, composed of 3–9 planes at 8–10 *µ*m z-spacing, was first registered to a larger anatomical imaging volume of the same animal acquired at the end of the experiment (1–2 *µ*m z-spacing) using affine and warp transformations. Depending on the transgenic line, the larger anatomical volume was then registered to the ZBB atlas using one of these strategies: For Tg(*elavl3*:H2B-GCaMP6s) larvae, we used a Tg(*elavl3*:H2B-GCaMP6s) reference volume (mean of 3 larvae) registered to ZBB using the ZBB Tg(*elavl3*:H2B-RFP) volume. For *u523* datasets, we used the ZBB Tg(*elavl3*:CAMPARI) volume. For *u524* datasets, we used a Tg(*u524*:Gal4FF; UAS:Kaede) reference volume (mean of 3 larvae) registered to ZBB via co-imaged Tg(*elavl3*:jRCaMP1a) using the ZBB Tg(*elavl3*:CAMPARI) volume. The transformations were concatenated to bring the functional imaging volume and associated soma centroids to ZBB space (calcium imaging stack → post-imaging stack → ZBB).
- PA-GFP imaging volumes were registered to the ZBB atlas using the ZBB Tg(*elavl3*:CAMPARI) volume and to the Mapzebrain atlas using a Tg(*elavl3*:jRCaMP1a) reference volume (mean of 3 larvae) registered to Mapzebrain using the Mapzebrain Tg(*gad1b*:GFP) volume.
- Imaging volumes of the retina were registered to a retinal Tg(*elavl3*:jRCaMP1a) reference volume (from 1 larva) via affine and warp transformations.

All registration steps were manually inspected for global and local alignment accuracy. Brain regions referred to in this paper correspond to the masks in ZBB (for somata) or Mapzebrain (for neurites), with the exception of Pt7 (a.k.a. AF7-pretectum), which is defined in ^18^.

### Optogenetics

Larval zebrafish were mounted as for calcium imaging (see above) at 5 or 6 dpf and allowed to recover overnight before testing at 6 or 7 dpf. Patterned illumination was delivered using a custom-built rig based on a digital micromirror device (DMD). The DMD (Texas Instruments DLP LightCrafter 6500 0.65 1080p) comprised 1920 × 1080 micro-mirrors (7.56 × 7.56 *µ*m) and was illuminated via an optical module (ViALUX STAR-065 CORE s600) connected to a liquid light-guide using either a 470 nm LED (Mightex BLS-LCS-0470-50-22) or a 6500 K white LED (Mightex BLS-LCS-6500-33-22) equipped with a 560/14 nm bandpass filter (Semrock FF01-560/14-25). LED sources were combined via a dichroic beam combiner (Mightex LCS-BC25-0495). The 470 nm excitation was used for activating CoChR-expressing neurons while 560 nm light was used to excite tdTomato in order to define target regions and acquire anatomical reference images. A relayed image of the DMD was projected onto the sample plane of the objective (Nikon CFI75 LWD 16× 0.8 NA; FF493/774-Di01-25×36 dichroic) such that individual micromirrors were 0.5 × 0.5 *µ*m at sample. For fluorescence detection, we used a CMOS camera (Hamamatsu ORCA-Flash4.0 V3) equipped with a 512–630 nm bandpass filter (Semrock FF01-512/630-25). To monitor behaviour, larvae were imaged at 400 Hz under 850 nm illumination using a sub-stage camera (Point Grey FL3-U3-13Y3M-C). Instrument control was achieved using custom routines written in LabVIEW (National Instruments).

Optogenetic stimulation comprised continuous illumination for 1 s at 2–10 mW/mm^2^ targeted to one or more 36 × 36 *µ*m loci in Pt7 or OT or control regions in parts of the larva lacking opsin expression. A minimum of three stimulation trials were performed for each target region.

### Photo-activation of PA-GFP

Larvae homozygous for the Tg(*Cau.Tuba1*:c3PA-GFP) and Tg(*elavl3*:jRCaMP1a) transgenes were anaesthetised and mounted in 2% low-melting temperature agarose at 5 dpf. Photo-activation were performed using same custom-built 2-photon microscope described above. Photo-activation loci in AF7, Pt7, OT or ventro-medial hindbrain were identified by imaging jRCaMP1a at 1040 nm. Next, PA-GFP was photo-activated by scanning a 9 × 9 × 40 *µ*m volume (790 nm, 5–10 mW at sample, 4–5 min per z-plane, 5–8 *µ*m plane spacing). Larvae were then unmounted and allowed to recover. Photo-labelled samples were imaged at 6–7 dpf by acquiring an image stack (1040 nm, 1200 × 1200 px, 0.38 *µ*m/px, 200 − 250 *µ*m z-extent, 2 *µ*m plane spacing) that encompassed a large portion of the forebrain, midbrain and hindbrain, or the retina contralateral to the photo-activation site. Soma locations were manually identified using the Multi-point tool in Fiji and neurites were traced using the Simple Neurite Tracer plugin in Fiji ^75^.

### Analysis of photolabelled neurites

We applied a clustering routine to PA-GFP-labelled neurites as well as previously published single neuron morphologies. First, a feature vector was generated for each neurite/neuron. Each feature vector consisted of the following components: (a) Minimum distances to each of 79 Mapzebrain regions in left and right hemispheres; (b) Number of neurite segments in each of the Mapzebrain regions in left and right hemispheres; (c) Minimum distance from soma to each of the Mapzebrain regions (only for single neuron morphologies as the exact soma location was not available for PA-GFP tracings); (d) Neurite path length. The full set of feature vectors was assembled into a matrix and each component was normalised by z-scoring. Single neuron morphologies and PA-GFP neurites were then clustered seperately using a hierarchical agglomerative clustering procedure with a correlation distance metric ^14^. To compare clusters, correlation coefficients were computed between mean feature vectors (excluding soma distance components).

### Laser axotomies

To enable the targeting and ablation of axon tracts, PA-GFP was photo-activated bilaterally (in pretectum for ablation of the crossed pretectobulbar pathway and in a-, m- and pOT for ablation of the ipsilateral tectobulbar pathway) at 5 dpf and laser axotomy was subsequently performed at 6 dpf. Larvae were anaesthetised using MS222 and mounted in 3% low-melting temperature agarose. Axotomy were performed using the same 2-photon microscope described above. Spiral scans (140 ms, 800 nm, 150–200 mW at sample) were performed at 5–10 locations centred on axonal tracts labelled by PA-GFP for crossed bulbar projections or tdTomato for uncrossed bulbar projections. Larvae were then allowed to recover overnight. Pre- and post-ablation image volumes were acquired at 920 nm (800 × 800 px, 0.38 *µ*m/px, 80 − 100 *µ*m z-extent, 2 *µ*m plane spacing).

### Statistical Analysis

All model fitting and statistical analyses were performed in MATLAB. Types of statistical test and *N* are reported in the text or figure legends. All tests were two-tailed and were chosen after data were tested for normality and homoscedasticity.

## Figures

**Figure S1:**
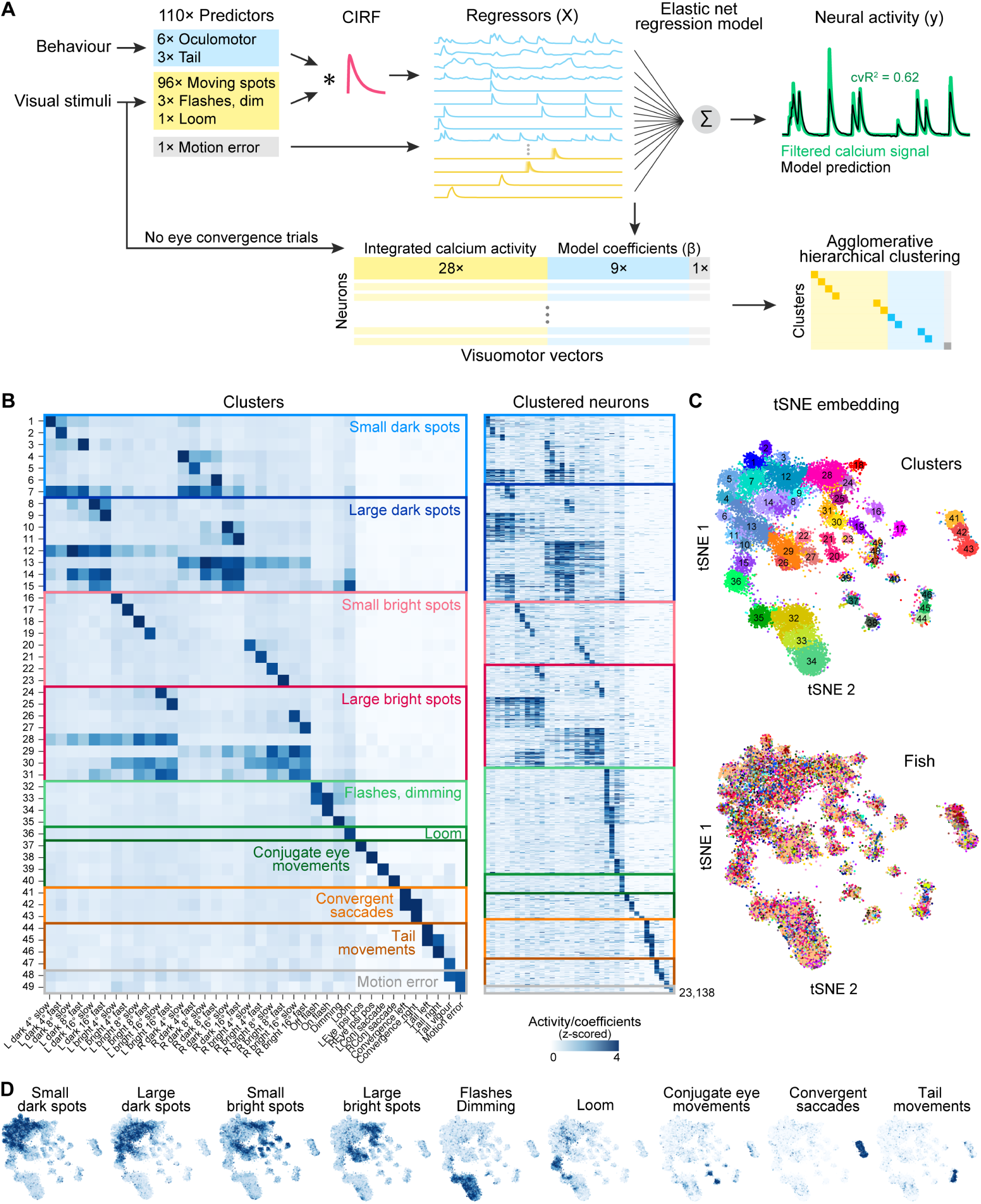
Functional clusters. (A) Generation of visuomotor vectors (VMVs) for individual neurons. CIRF, calcium impulse response function. cv*R*^2^, cross-validated goodness of fit. See Methods for details. (B) Cluster centroids (left) and VMVs of all neurons (right, 23,138 cells). Colored boxes and labels indicate broad functional categories. (C) t-distributed stochastic neighbour embedding (t-SNE) of VMVs of clustered cells. Top, coded by cluster identity; bottom, coded by fish identity. (D) t-SNE embedding coded by the indicated VMV coefficients. For categories having multiple coefficients (e.g. small dark spots) the max coefficient value is used per cell.

**Figure S2:**
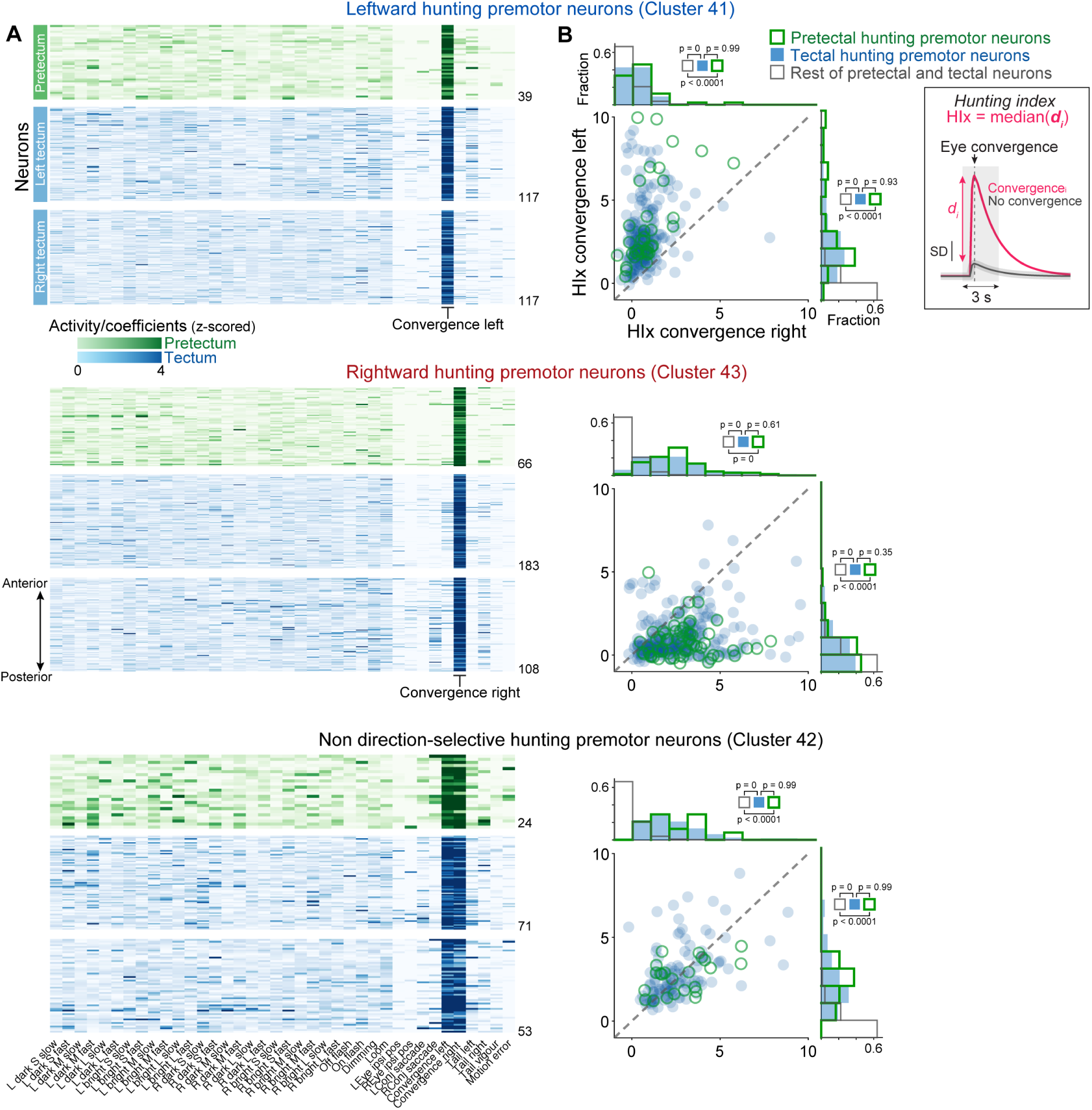
Activity of hunting premotor neurons. (A) VMVs for hunting premotor cells belonging to cluster 41 (left tuned, top), cluster 43 (right tuned, middle) and cluster 42 (direction agnostic, bottom). Within each cluster, cells are sorted according to anterior-posterior location. Number cells indicated on the right. Premotor cells display high VMV coefficients for convergent saccades but minimal activity to visual stimuli or non-hunting tail movements. (B) Hunting Index (HIx) ^18^, which quantifies activity modulation triggered on hunting initiation (detected by convergent saccades; see Methods and inset box). Plots show HIx values for each cell for left- and right-directed hunting events. Marginal distributions and Tukey HSD-adjusted *p*-values of multiple comparison tests following three-sample Kruskal-Wallis tests are reported.

**Figure S3:**
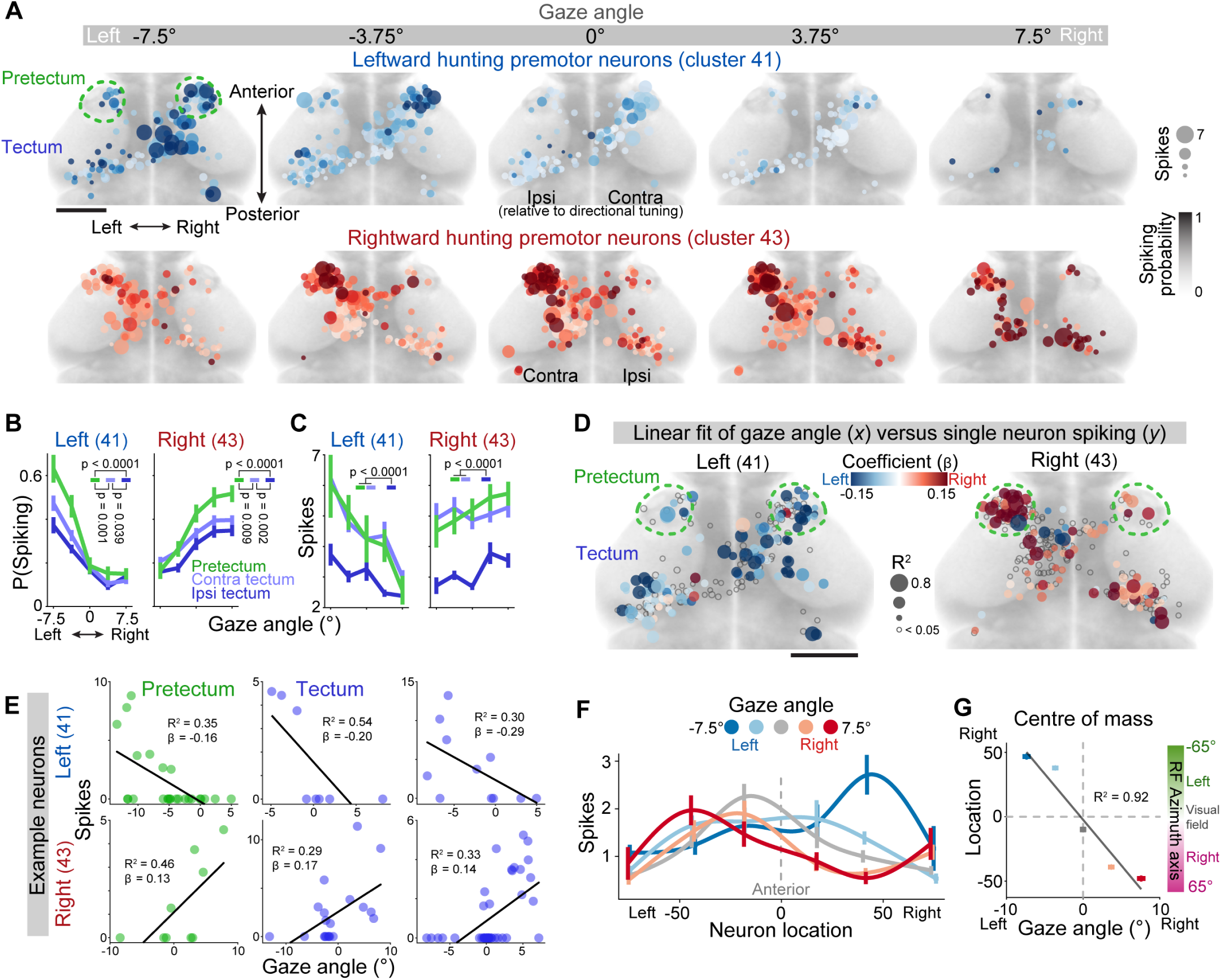
Premotor encoding of saccadic eye movements. (A) Activity maps for direction-selective premotor neurons (cluster 41, left-selective, *N* = 273 neurons; cluster 43, right-selective, *N* = 357 neurons from 22 larvae). OASIS-inferred spike probability (colour intensity) and spike counts (dot size) are shown for hunting manoeuvres binned by post-saccadic gaze (bin centres indicated at top). (B,C) Spike probability (B) and spike counts (C) versus gaze for premotor neurons in Pt7 (green), contralateral OT (light blue), and ipsilateral OT (dark blue), where ipsi/contralateral is with respect to the cells’ directional tuning. Mean ± SEM were computed across neurons, as for all the remaining plots (cluster 41: *N* = 39 pretectal neurons, *N* = 117 contra tectum, *N* = 117 ipsi tectum; cluster 43: *N* = 66 pretectal neurons, *N* = 183 contra tectum, *N* = 108 ipsi tectum). Tukey HSD-adjusted *p*-values for multiple comparison tests following two-way ANOVA. (D) Linear regression models relating single-cell spike counts (y) to post-saccadic gaze (x). Fit coefficients (*β*, colour) and goodness-of-fit values (*R*^2^, dot size). Left angles are negative, so neurons encoding leftward movement have negative coefficients. (E) Examples of single neuron linear regression fits. (F) Spatial distribution of premotor activity along the anterior-posterior axis of Pt7-OT. Each trace represents a different bin of post-saccadic gaze and shows spike counts across neurons (cluster 41 and 43 combined) binned by anatomical location. (G) Centre-of-mass of premotor activity for hunting manoeuvres binned by post-saccadic gaze. The azimuth axis obtained from the linear regression fit in Figure 1H is also reported on the right.

**Figure S4:**
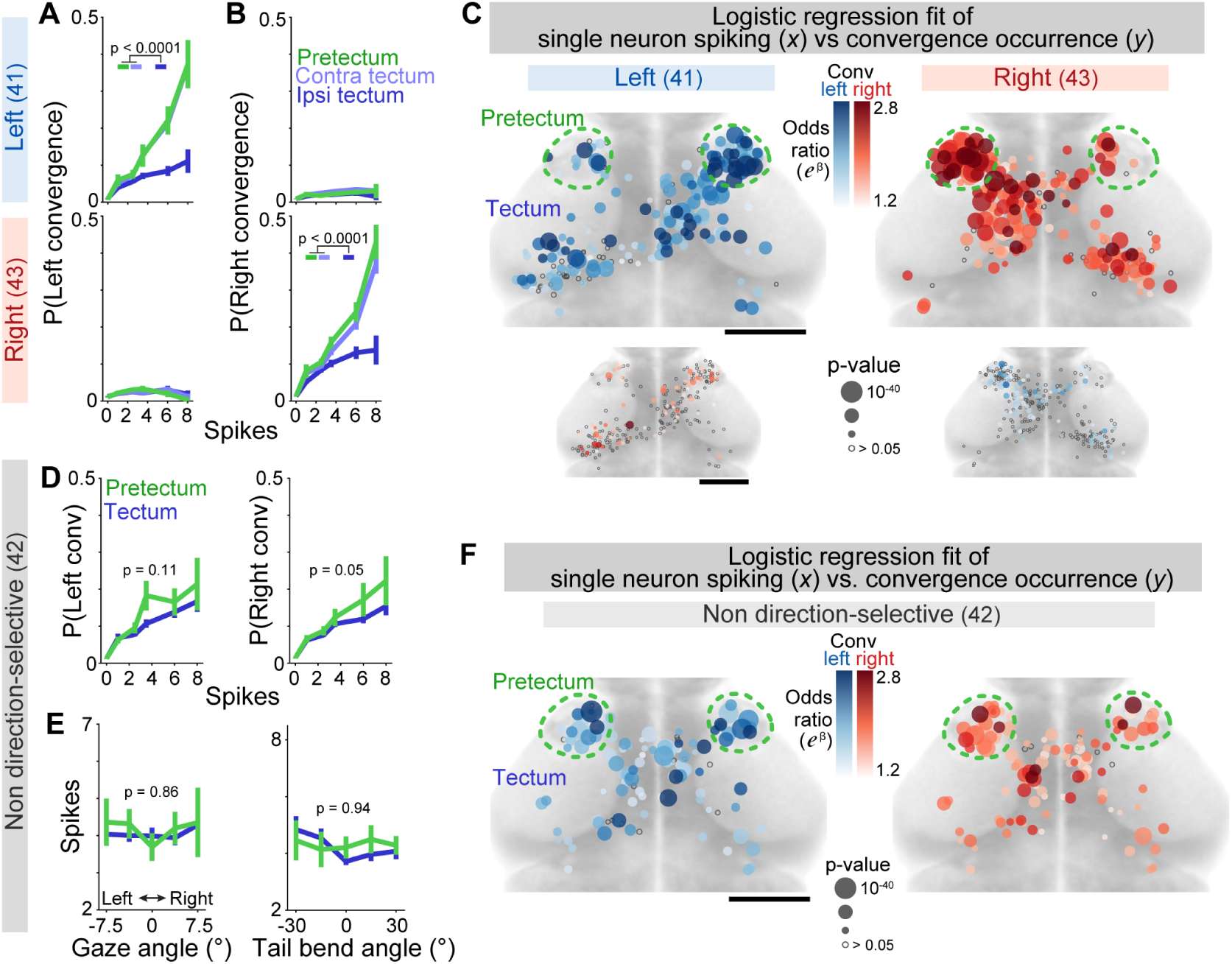
Premotor encoding of hunting response probability. (A,B) Probability of hunting initiation versus activity of direction-selective premotor clusters. Mean ±SEM were computed across neurons (cluster 41: *N* = 39 pretectal neurons, *N* = 117 contra tectum, *N* = 117 ipsi tectum; cluster 43: *N* = 66 pretectal neurons, *N* = 183 contra tectum, *N* = 108 ipsi tectum). Tukey HSD-adjusted *p*-values for multiple comparison tests following two-way ANOVA. (C) Logistic regression models for direction-selective premotor neurons. For each cell, we fit a model predicting hunting responses (y) from the number of OASIS-inferred spikes per imaging frame (x). Maps show odds ratios (*e^β^*, colour) and *p*-values (dot size). Scale bar: 100*µ*m. (D,E) Activity of non-direction-selective premotor neurons (cluster 42, *N* = 148 neurons from 15 larvae) scales with probability of hunting initiation (D) but not directionality of motor outputs (E). (F) Logistic regression models relating hunting initiation to activity for non-direction-selective premotor neurons, similar to (C).

**Figure S5:**
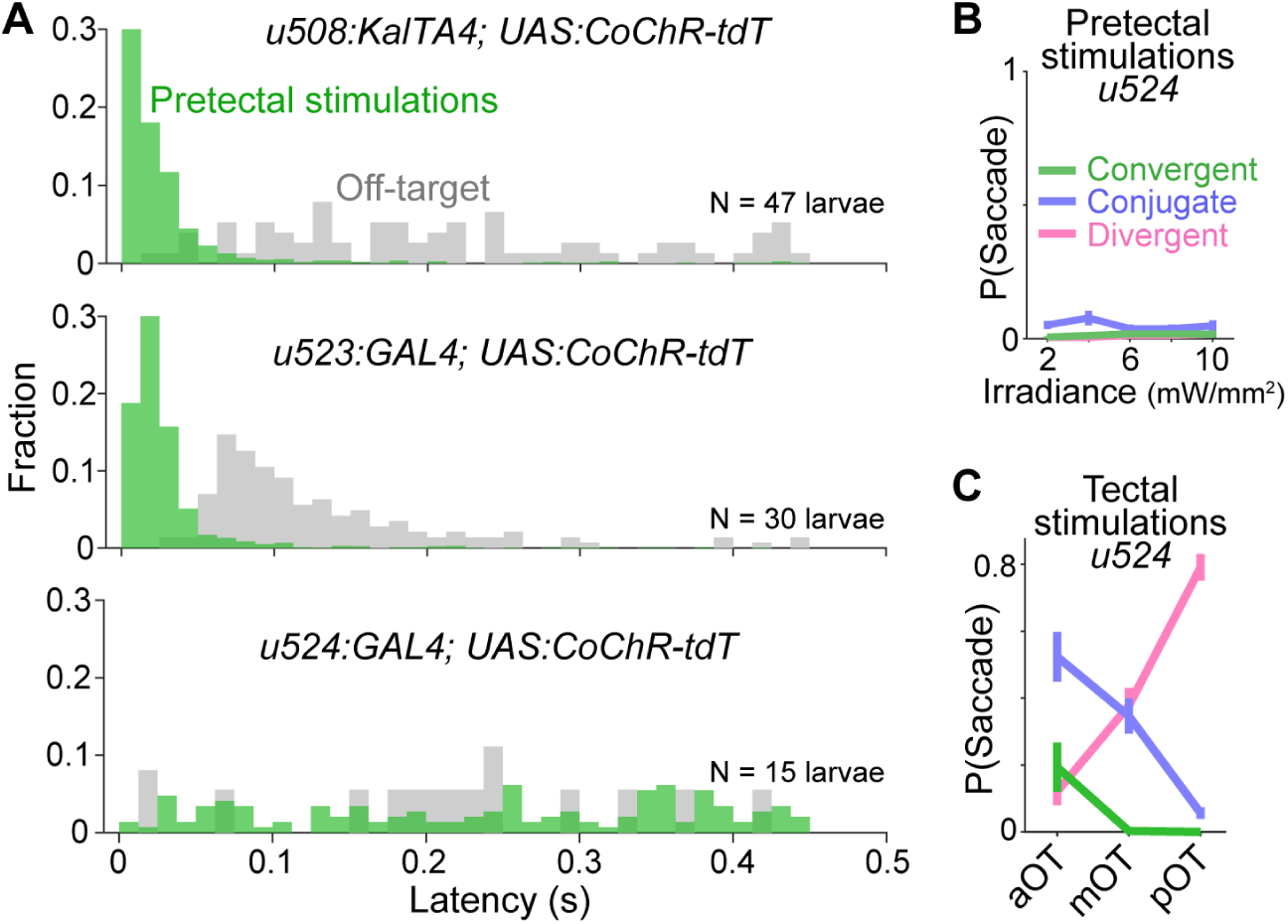
Optogenetic stimulation – additional data. (A) Response latency following stimulation of Pt7 (green) or off-target control regions (grey). (B) Pt7 stimulation in Tg(*u524*:GAL4;UAS:CoChR-tdTomato) larvae was not effective at inducing behaviour. *N* = 15 larvae. (C) OT stimulation in Tg(*u524*:GAL4;UAS:CoChR-tdTomato) larvae. Most optogenetically induced eye movements were conjugate or divergent saccades, rather than predatory convergent saccades. *N* = 20 larvae.

**Figure S6:**
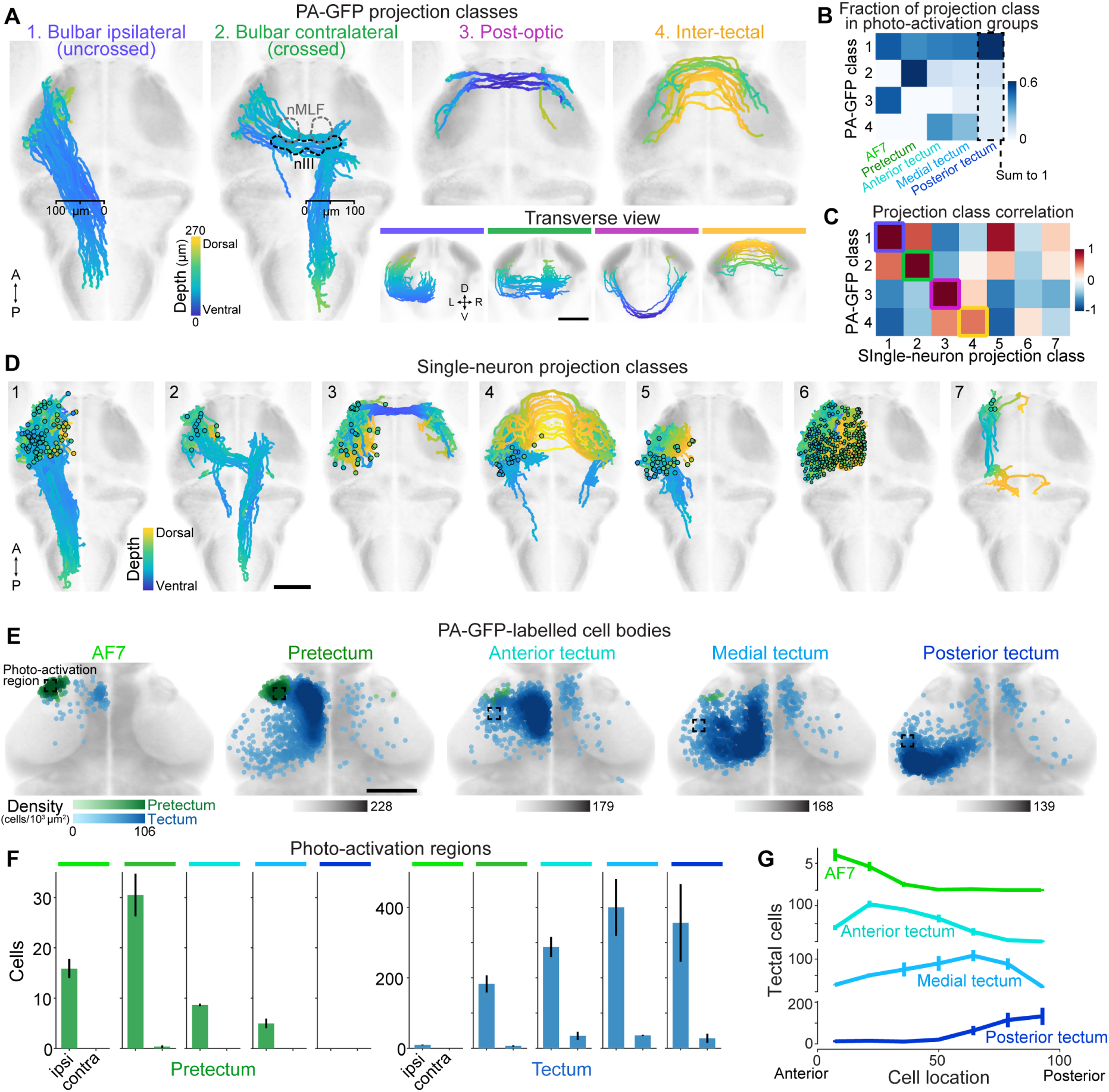
Projection classes and Pt7-OT reciprocal connections. (A) Projection classes obtained from morphological clustering of PA-GFP-labelled neurites (50 class 1 neurites, 20 class 2, 8 class 3, 19 class 4, from 35 larvae). Images are depth-coded (yellow dorsal, blue ventral) and transverse views are shown bottom right. Scale bar: 100 *µ*m; (B) Relative abundance of each projection class obtained from each photoactivation locus. Values in each column sum to one. (C) Correlation between mean anatomical feature vectors for PA-GFP projection classes and single-neuron projection classes. For each PA-GFP projection class, the most similar single-neuron projection class is indicated with a square. (D) Projection classes obtained from morphological clustering of single neurons with soma located in Pt7-OT, from ^6,18,19,33^. Scale bar: 100 *µ*m. (E) Labelled cell bodies in pretectum (green) and tectum (blue) following photoactivation at different loci in Pt7-OT. *N* = 7 larvae for each photoactivation locus. Scale bar: 100 *µ*m. (F) Counts of retrogradely labelled cells in ipsi- and contralateral pretectum (green) and tectum (blue) following photoactivation at each locus (indicated by coloured bars). Mean ± SEM was computed across fish. (G) Distribution of retrogradely labelled tectal cells across the anterior-posterior axis for each photoactivation locus. Equivalent data for pretectum is in Figure 6J.

**Figure S7:**
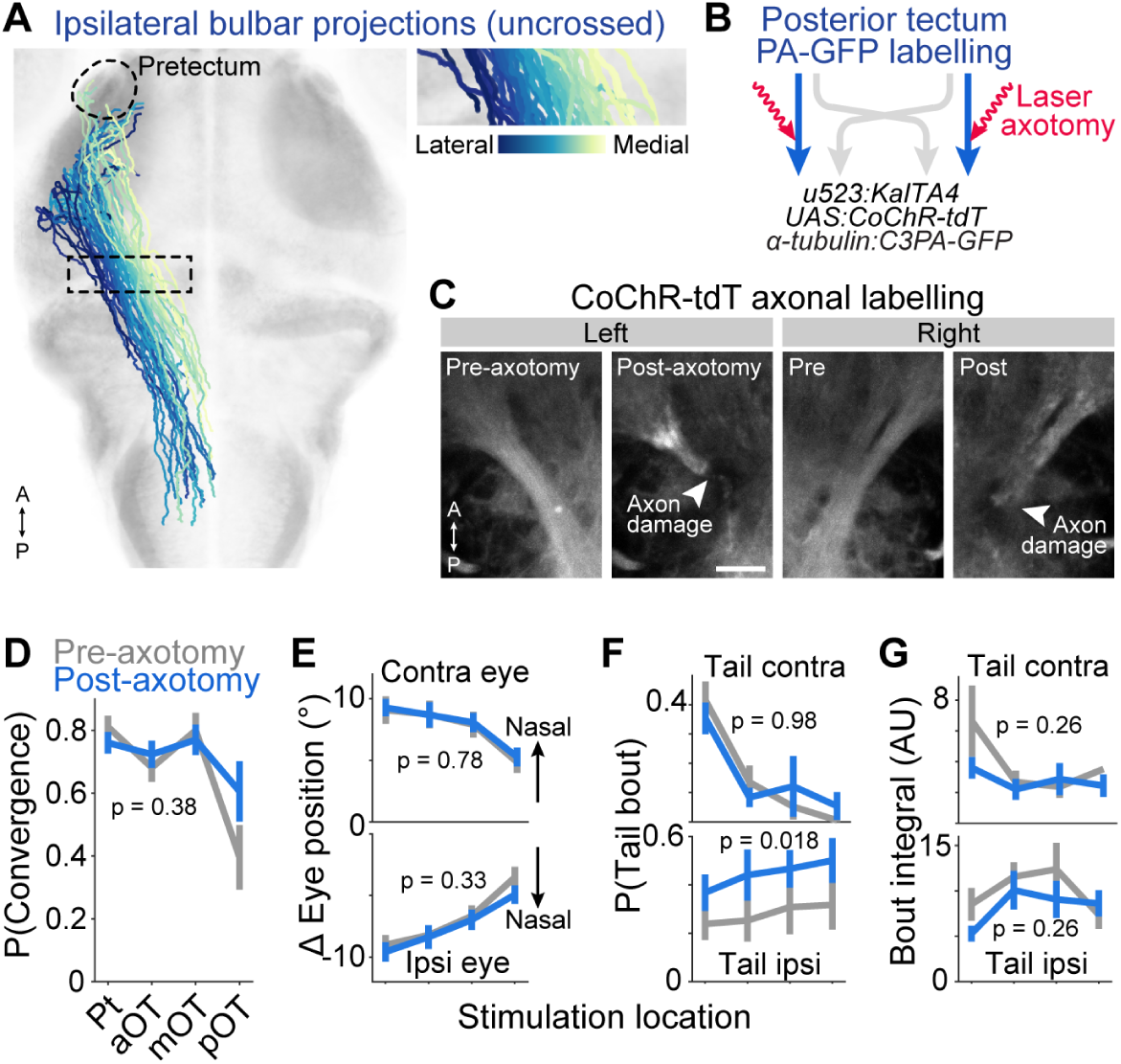
Axotomy of ipsilateral bulbar projections. (A) Uncrossed (type 1) bulbar projections colour-coded according to medio-lateral location of axons within the boxed region (enlarged in inset). Neurites originating from posterior tectum tended to occupy more lateral locations in the tract. (B) Schematic of laser axotomies. (C) Maximum intensity projections of regions in which CoChR-tdTomato-expressing bulbar projections originating from posterior tectum were targeted for axotomy, following bilateral PA-GFP photoactivation in pOT. Images prior to (6 dpf) and after (7 dpf) axotomy. Scale bar: 15 *µ*m. (D) Probability of evoking a convergent saccade versus stimulus location before (grey) and after (blue) laser axotomy. Mean ± SEM was computed across fish, as for all other plots (*N* = 7 larvae). (E) Amplitude of evoked eye movements. Ipsi and contra relative to stimulation site. (F) Probability of convergent saccade being paired with contraversive or ipsiversive tail movement. (G) Integrated tail bend angle for contraversive and ipsiversive tail movements. In D-G, *p*-values from two-way ANOVA comparing pre- vs post-axotomy.

## References

1. Isa, T., Marquez-Legorreta, E., Grillner, S., and Scott, E.K. (2021). The tectum/superior colliculus as the vertebrate solution for spatial sensory integration and action. Curr Biol 31, R741–R762.

2. Gandhi, N.J. and Katnani, H.A. (2011). Motor functions of the superior colliculus. Annual review of neuroscience 34, 205–31.

3. Dean, P., Redgrave, P., Sahibzada, N., and Tsuji, K. (1986). Head and body movements produced by electrical stimulation of superior colliculus in rats: effects of interruption of crossed tectoreticulospinal pathway. Neuroscience 19, 367–80.

4. Dean, P., Redgrave, P., and Westby, G.W. (1989). Event or emergency? Two response systems in the mammalian superior colliculus. Trends Neurosci 12, 137–47.

5. Kostyk, S.K. and Grobstein, P. (1987). Neuronal organization underlying visually elicited prey orienting in the frog–III. Evidence for the existence of an uncrossed descending tectofugal pathway. Neuroscience 21, 83–96.

6. Helmbrecht, T.O., Dal Maschio, M., Donovan, J.C., Koutsouli, S., and Baier, H. (2018). Topography of a Visuomotor Transformation. Neuron 100, 1429–1445.e4.

7. Borla, M.A., Palecek, B., Budick, S., and O’Malley, D.M. (2002). Prey capture by larval zebrafish: evidence for fine axial motor control. Brain Behav Evol 60, 207–29.

8. McElligott, M.B. and O’Malley, D.M. (2005). Prey tracking by larval zebrafish: axial kinematics and visual control. Brain Behav Evol 66, 177–96.

9. Bianco, I.H., Kampff, A.R., and Engert, F. (2011). Prey capture behavior evoked by simple visual stimuli in larval zebrafish. Front Syst Neurosci 5, 101.

10. Trivedi, C.A. and Bollmann, J.H. (2013). Visually driven chaining of elementary swim patterns into a goal-directed motor sequence: a virtual reality study of zebrafish prey capture. Front Neural Circuits 7, 86.

11. Bolton, A.D., Haesemeyer, M., Jordi, J., Schaechtle, U., Saad, F.A., Mansinghka, V.K., Tenenbaum, J.B., and Engert, F. (2019). Elements of a stochastic 3D prediction engine in larval zebrafish prey capture. Elife 8.

12. Mearns, D.S., Donovan, J.C., Fernandes, A.M., Semmelhack, J.L., and Baier, H. (2020). Deconstructing Hunting Behavior Reveals a Tightly Coupled Stimulus-Response Loop. Curr Biol 30, 54–69.e9.

13. Johnson, R.E., Linderman, S., Panier, T., Wee, C.L., Song, E., Herrera, K.J., Miller, A., and Engert, F. (2020). Probabilistic Models of Larval Zebrafish Behavior Reveal Structure on Many Scales. Curr Biol 30, 70–82.e4.

14. Bianco, I.H. and Engert, F. (2015). Visuomotor transformations underlying hunting behavior in zebrafish. Curr Biol 25, 831–46.

15. Gahtan, E., Tanger, P., and Baier, H. (2005). Visual prey capture in larval zebrafish is controlled by identified reticulospinal neurons downstream of the tectum. J Neurosci 25, 9294–303.

16. Niell, C.M. and Smith, S.J. (2005). Functional imaging reveals rapid development of visual response properties in the zebrafish tectum. Neuron 45, 941–51.

17. Preuss, S.J., Trivedi, C.A., vom Berg-Maurer, C.M., Ryu, S., and Bollmann, J.H. (2014). Classification of object size in retinotectal microcircuits. Current Biology 24, 2376–2385.

18. Antinucci, P., Folgueira, M., and Bianco, I.H. (2019). Pretectal neurons control hunting behaviour. Elife 8.

19. Förster, D., Helmbrecht, T.O., Mearns, D.S., Jordan, L., Mokayes, N., and Baier, H. (2020). Retinotectal circuitry of larval zebrafish is adapted to detection and pursuit of prey. Elife 9, e58596.

20. Zylbertal, A. and Bianco, I.H. (2023). Recurrent network interactions explain tectal response variability and experience-dependent behavior. Elife 12.

21. Fajardo, O., Zhu, P., and Friedrich, R.W. (2013). Control of a specific motor program by a small brain area in zebrafish. Front Neural Circuits 7, 67.

22. Semmelhack, J.L., Donovan, J.C., Thiele, T.R., Kuehn, E., Laurell, E., and Baier, H. (2014). A dedicated visual pathway for prey detection in larval zebrafish. Elife 3, e04878.

23. Dowell, C.K., Lau, J.Y.N., Antinucci, P., and Bianco, I.H. (2024). Kinematically distinct saccades are used in a context-dependent manner by larval zebrafish. Curr Biol, S0960–9822(24)01084–4.

24. Muto, A., Lal, P., Ailani, D., Abe, G., Itoh, M., and Kawakami, K. (2017). Activation of the hypothalamic feeding centre upon visual prey detection. Nat Commun 8, 15029.

25. Henriques, P.M., Rahman, N., Jackson, S.E., and Bianco, I.H. (2019). Nucleus Isthmi Is Required to Sustain Target Pursuit during Visually Guided Prey-Catching. Curr Biol 29, 1771–1786.e5.

26. Marquart, G.D., Tabor, K.M., Brown, M., Strykowski, J.L., Varshney, G.K., LaFave, M.C., Mueller, T., Burgess, S.M., Higashijima, S.i., and Burgess, H.A. (2015). A 3D searchable database of transgenic zebrafish Gal4 and Cre lines for functional neuroanatomy studies. Frontiers in neural circuits 9, 78.

27. Marquart, G.D., Tabor, K.M., Horstick, E.J., Brown, M., Geoca, A.K., Polys, N.F., Nogare, D.D., and Burgess, H.A. (2017). High-precision registration between zebrafish brain atlases using symmetric diffeomorphic normalization. GigaScience 6, gix056.

28. Friedrich, J., Zhou, P., and Paninski, L. (2017). Fast online deconvolution of calcium imaging data. PLoS Comput Biol 13, e1005423.

29. Robles, E., Laurell, E., and Baier, H. (2014). The retinal projectome reveals brain-area-specific visual representations generated by ganglion cell diversity. Curr Biol 24, 2085–96.

30. Schmitt, E.A. and Dowling, J.E. (1999). Early retinal development in the zebrafish, Danio rerio: light and electron microscopic analyses. J Comp Neurol 404, 515–36.

31. Yoshimatsu, T., Schröder, C., Nevala, N.E., Berens, P., and Baden, T. (2020). Fovea-like Photoreceptor Specializations Underlie Single UV Cone Driven Prey-Capture Behavior in Zebrafish. Neuron 107, 320–337.e6.

32. Antinucci, P., Dumitrescu, A., Deleuze, C., Morley, H.J., Leung, K., Hagley, T., Kubo, F., Baier, H., Bianco, I.H., and Wyart, C. (2020). A calibrated optogenetic toolbox of stable zebrafish opsin lines. Elife 9.

33. Kunst, M., Laurell, E., Mokayes, N., Kramer, A., Kubo, F., Fernandes, A.M., Förster, D., Dal Maschio, M., and Baier, H. (2019). A Cellular-Resolution Atlas of the Larval Zebrafish Brain. Neuron 103, 21–38.e5.

34. Robinson, D.A. (1972). Eye movements evoked by collicular stimulation in the alert monkey. Vision Res 12, 1795–808.

35. Schiller, P.H. and Stryker, M. (1972). Single-unit recording and stimulation in superior colliculus of the alert rhesus monkey. J Neurophysiol 35, 915–24.

36. Lee, C., Rohrer, W.H., and Sparks, D.L. (1988). Population coding of saccadic eye movements by neurons in the superior colliculus. Nature 332, 357–60.

37. Sparks, D.L. and Mays, L.E. (1990). Signal transformations required for the generation of saccadic eye movements. Annu Rev Neurosci 13, 309–36.

38. Munoz, D.P., Guitton, D., and Pélisson, D. (1991). Control of orienting gaze shifts by the tectoreticulospinal system in the head-free cat. III. Spatiotemporal characteristics of phasic motor discharges. J Neurophysiol 66, 1642–66.

39. Paré, M., Crommelinck, M., and Guitton, D. (1994). Gaze shifts evoked by stimulation of the superior colliculus in the head-free cat conform to the motor map but also depend on stimulus strength and fixation activity. Exp Brain Res 101, 123–39.

40. Freedman, E.G., Stanford, T.R., and Sparks, D.L. (1996). Combined eye-head gaze shifts produced by electrical stimulation of the superior colliculus in rhesus monkeys. J Neurophysiol 76, 927–52.

41. Stanford, T.R., Freedman, E.G., and Sparks, D.L. (1996). Site and parameters of microstimulation: evidence for independent effects on the properties of saccades evoked from the primate superior colliculus. J Neurophysiol 76, 3360–81.

42. González-Rueda, A., Jensen, K., Noormandipour, M., de Malmazet, D., Wilson, J., Ciabatti, E., Kim, J., Williams, E., Poort, J., Hennequin, G., et al. (2024). Kinetic features dictate sensorimotor alignment in the superior colliculus. Nature 631, 378–385.

43. Moschovakis, A.K., Kitama, T., Dalezios, Y., Petit, J., Brandi, A.M., and Grantyn, A.A. (1998). An anatomical substrate for the spatiotemporal transformation. J Neurosci 18, 10219–29.

44. Herrero, L., Rodriguez, F., Salas, C., and Torres, B. (1998). Tail and eye movements evoked by electrical microstimulation of the optic tectum in goldfish. Exp Brain Res 120, 291–305.

45. Sahibzada, N., Dean, P., and Redgrave, P. (1986). Movements resembling orientation or avoidance elicited by electrical stimulation of the superior colliculus in rats. J Neurosci 6, 723–33.

46. du Lac, S. and Knudsen, E.I. (1990). Neural maps of head movement vector and speed in the optic tectum of the barn owl. J Neurophysiol 63, 131–46.

47. Zahler, S.H., Taylor, D.E., Wright, B.S., Wong, J.Y., Shvareva, V.A., Park, Y.A., and Feinberg, E.H. (2023). Hindbrain modules differentially transform activity of single collicular neurons to coordinate movements. Cell 186, 3062–3078.e20.

48. Ellard, C.G. and Goodale, M.A. (1988). A functional analysis of the collicular output pathways: a dissociation of deficits following lesions of the dorsal tegmental decussation and the ipsilateral collicular efferent bundle in the Mongolian gerbil. Exp Brain Res 71, 307–19.

49. Keay, K., Westby, G.W., Frankland, P., Dean, P., and Redgrave, P. (1990). Organization of the crossed tecto-reticulo-spinal projection in rat–II. Electrophysiological evidence for separate output channels to the periabducens area and caudal medulla. Neuroscience 37, 585–601.

50. Redgrave, P., Dean, P., and Westby, G.W. (1990). Organization of the crossed tecto-reticulo-spinal projection in rat–I. Anatomical evidence for separate output channels to the periabducens area and caudal medulla. Neuroscience 37, 571–84.

51. Ingle, D.J. (1983). Brain Mechanisms of Visual Localization by Frogs and Toads in Advances in Vertebrate Neuroethology (Springer US).

52. Winterkorn, J.M.S. and Meikle, T.H. (1980). Lesions of the tectospinal tract of the cat do not produce compulsive circling. Brain Research 190, 597–600.

53. Masino, T. and Grobstein, P. (1989). The organization of descending tectofugal pathways underlying orienting in the frog, Rana pipiens. I. Lateralization, parcellation, and an intermediate spatial representation. Exp Brain Res 75, 227–44.

54. Masino, T. and Grobstein, P. (1989). The organization of descending tectofugal pathways underlying orienting in the frog, Rana pipiens. II. Evidence for the involvement of a tecto-tegmento-spinal pathway. Exp Brain Res 75, 245–64.

55. King, J.R. and Comer, C.M. (1996). Visually elicited turning behavior in Rana pipiens: comparative organization and neural control of escape and prey capture. J Comp Physiol A 178, 293–305.

56. Isa, K., Sooksawate, T., Kobayashi, K., Kobayashi, K., Redgrave, P., and Isa, T. (2020). Dissecting the Tectal Output Channels for Orienting and Defense Responses. eNeuro 7, ENEURO.0271–20.2020.

57. Tian, G., Lam, T.K.C., Yan, G., He, Y., Khan, B., Qu, J.Y., and Semmelhack, J.L. (2025). Binocular integration of prey stimuli in the zebrafish visual system. Curr Biol 35, 3228–3240.e5.

58. Gebhardt, C., Auer, T.O., Henriques, P.M., Rajan, G., Duroure, K., Bianco, I.H., and Del Bene, F. (2019). An interhemispheric neural circuit allowing binocular integration in the optic tectum. Nat Commun 10, 5471.

59. Lister, J.A., Robertson, C.P., Lepage, T., Johnson, S.L., and Raible, D.W. (1999). nacre encodes a zebrafish microphthalmia-related protein that regulates neural-crest-derived pigment cell fate. Development 126, 3757– 67.

60. Freeman, J., Vladimirov, N., Kawashima, T., Mu, Y., Sofroniew, N.J., Bennett, D.V., Rosen, J., Yang, C.T., Looger, L.L., and Ahrens, M.B. (2014). Mapping brain activity at scale with cluster computing. Nat Methods 11, 941–50.

61. Dowell, C.K., Hawkins, T., and Bianco, I.H. (2025). Subsets of extraocular motoneurons produce kinematically distinct saccades during hunting and exploration. Curr Biol 35, 554–573.e6.

62. Bianco, I.H., Ma, L.H., Schoppik, D., Robson, D.N., Orger, M.B., Beck, J.C., Li, J.M., Schier, A.F., Engert, F., and Baker, R. (2012). The tangential nucleus controls a gravito-inertial vestibulo-ocular reflex. Curr Biol 22, 1285–95.

63. Dunn, T.W., Mu, Y., Narayan, S., Randlett, O., Naumann, E.A., Yang, C.T., Schier, A.F., Freeman, J., Engert, F., and Ahrens, M.B. (2016). Brain-wide mapping of neural activity controlling zebrafish exploratory locomotion. Elife 5, e12741.

64. Davison, J.M., Akitake, C.M., Goll, M.G., Rhee, J.M., Gosse, N., Baier, H., Halpern, M.E., Leach, S.D., and Parsons, M.J. (2007). Transactivation from Gal4-VP16 transgenic insertions for tissue-specific cell labeling and ablation in zebrafish. Dev Biol 304, 811–24.

65. Brainard, D.H. (1997). The Psychophysics Toolbox. Spat Vis 10, 433–6.

66. Sun, H. and Frost, B.J. (1998). Computation of different optical variables of looming objects in pigeon nucleus rotundus neurons. Nature neuroscience 1, 296.

67. Dunn, T.W., Gebhardt, C., Naumann, E.A., Riegler, C., Ahrens, M.B., Engert, F., and Del Bene, F. (2016). Neural Circuits Underlying Visually Evoked Escapes in Larval Zebrafish. Neuron 89, 613–28.

68. Lau, J.Y.N., Fitzgerald, J.E., and Bianco, I.H. (2025). Supraspinal commands have a modular organization that is behavioral context specific. Curr Biol, S0960–9822(25)01003–6.

69. Kawashima, T., Zwart, M.F., Yang, C.T., Mensh, B.D., and Ahrens, M.B. (2016). The serotonergic system tracks the outcomes of actions to mediate short-term motor learning. Cell 167, 933–946.

70. Pachitariu, M., Stringer, C., Dipoppa, M., Schroder, S.L., Rossi, F., Dalgleish, H., Carandini, M., and Harris, K.D. (2017). Suite2p: beyond 10,000 neurons with standard two-photon microscopy. biorxiv .

71. Zou, H. and Hastie, T. (2005). Regularization and variable selection via the elastic net. J. R. Stat. Soc. Ser. B-Stat. Methodol. 67, 301–320. URL http://ws.isiknowledge.com/cps/openurl/service?url_ver=Z39.88-2004&rft_id=info:ut/WOS:000227498200007.

72. Qian, J., Hastie, T., Friedman, J., Tibshirani, R., and Simon, N. (2013). Glmnet for Matlab. URL http://hastie.su.domains/glmnet_matlab/.

73. Sajovic, P. and Levinthal, C. (1982). Visual cells of zebrafish optic tectum: mapping with small spots. Neuroscience 7, 2407–26.

74. Avants, B.B., Tustison, N.J., Song, G., Cook, P.A., Klein, A., and Gee, J.C. (2011). A reproducible evaluation of ANTs similarity metric performance in brain image registration. Neuroimage 54, 2033–2044.

75. Longair, M.H., Baker, D.A., and Armstrong, J.D. (2011). Simple Neurite Tracer: open source software for reconstruction, visualization and analysis of neuronal processes. Bioinformatics 27, 2453–2454.

76. Easter Jr, S.S. and Nicola, G.N. (1996). The development of vision in the zebrafish (Danio rerio). Developmental biology 180, 646–663.

